# Prediction of combination therapies based on topological modeling of the immune signaling network in Multiple Sclerosis

**DOI:** 10.1101/541458

**Authors:** Marti Bernardo-Faura, Melanie Rinas, Jakob Wirbel, Inna Pertsovskaya, Vicky Pliaka, Dimitris E Messinis, Gemma Vila, Theodore Sakellaropoulos, Wolfgang Faigle, Pernilla Stridh, Janina R. Behrens, Tomas Olsson, Roland Martin, Friedemann Paul, Leonidas G Alexopoulos, Pablo Villoslada, Julio Saez-Rodriguez

## Abstract

Signal transduction deregulation is a hallmark of many complex diseases, including Multiple Sclerosis (MS). Here, we performed *ex vivo* multiplexed phosphoproteomic assays upon perturbations with multiple drugs and ligands in primary immune cells from 169 MS patients and matched healthy controls. Patients were either untreated or treated with fingolimod, natalizumab, interferon-β, glatiramer acetate or the experimental therapy epigallocatechin gallate (EGCG). We generated for each donor a dynamic logic model by fitting a bespoke literature-derived network of MS-related pathways to the perturbation data. Analysis of the models uncovered features of healthy-, disease- and drug-specific signaling networks. We developed an approach based on network topology to identify deregulated interactions whose activity could be reverted to a “healthy-like” status by combination therapy. We predicted several combinations with approved MS drugs. Specifically, TAK1 kinase, involved in TGF-β, Toll-like receptor, B-cell receptor and response to inflammation pathways were found to be highly deregulated and co-druggable with all MS drugs studied. One of these predicted combinations, fingolimod with a TAK1 inhibitor, was validated in an animal model of MS. Our approach based on donor-specific signaling networks enables prediction of targets for combination therapy for MS and other complex diseases.

**Significance statement:** Multiple Sclerosis (MS) is a major health problem, leading to significant disability and patient suffering. Although chronic activation of the immune system is a hallmark of the disease, its pathogenesis is poorly understood. Further, current treatments only ameliorate the disease and may produce severe side effects.

Here, we applied a network-based modeling approach based on phosphoproteomic data upon perturbation with ligands and drugs of healthy donors and MS patients to create donor-specific models. The models uncover the differential activation in signaling wiring between healthy donors, untreated patients and those under different treatments. Further, based in the patient-specific networks, a new approach identifies drug combinations to revert signaling to a healthy-like state.

**One sentence summary:** A new approach to predict combination therapies based on modeling signaling networks using phosphoproteomics from Multiple Sclerosis patients identifies deregulated pathways and new drug combinations.

## Introduction

The signal transduction machinery is frequently affected by perturbations induced by complex diseases. Hence, treatment of diseases such as cancer, cardiovascular, immunological or brain diseases is nowadays largely attempted by modulating different molecular cascades involved in the disease to stop its progression. As such, kinases involved in signaling processes have evolved as primary targets for many diseases (1). Further, there has been modest progress with treatments based on single drugs. Combining several drugs targeting different pathways promises more effective modulation of the pathogenic process (2, 3). However, development of combination therapies is hampered by the often incomplete understanding on how their effect propagates through complex signaling networks, with crosstalk between the pathways influenced by each therapy (4, 5). As an additional level of complication, disease heterogeneity hinders predicting how a specific combination therapy could be translated into the clinic. Last, the combinatorial nature of such studies in terms of number of targets, drugs, doses, and therapeutic regimens implies a large number of experiments and associated costs, preventing a complete analysis for all alternatives. As a result, the full potential of combination therapies has not been fully developed yet.

Systems biology, and more specifically modeling of signaling pathways applied to drug discovery, may provide a new path to approach this question (2, 6). Mechanistic understanding at the network level offers integrated insights about the cellular responses to environmental changes (7) and drug effects, yielding a significant understanding of the signaling cascades derived from decades of research in this field (8–10). Mathematical modeling of signaling networks has been used to unravel signaling mechanisms, discover drug targets and disease mediators such as cell surface receptors or intracellular molecules by training those models to experimental *in vitro* measurements of key pathway components using inhibitors and activators (4, 11, 12). Modeling-based studies may be the key to characterize the effect of drug combinations at the molecular level and allow us to predict both efficacy and reduction of off-target effects (5, 11, 13).

Multiple Sclerosis (MS) is an autoimmune disease, in which the immune system is chronically activated and damages the central nervous system (CNS) (14). The involvement of immune system deregulation in MS is shown by altered phenotype and activity of blood lymphocytes and monocytes (14), as well as by association with genetic polymorphisms of immune genes (15). At present, there are fifteen FDA-approved immunomodulatory drugs, and many others are in late stage clinical development. Most of these control the inflammatory activity in patients with MS to a certain degree, though with known unwanted effects at the signaling level. Furthermore, many unmet medical needs remain in the attempt of achieving control of the disease including more effective therapies, a good safety profile and neuroprotective or regenerative treatments. Drug combinations are considered as a promising strategy to overcome some of these limitations, and in cancer combination therapies they are well established (16). However, predicting which patients would benefit the most from a certain combination therapy remains as an unresolved challenge (17–20).

In this study we present a systems medicine approach aimed to (i) characterize the signaling pathways in primary immune cells obtained from blood of MS patients and healthy controls, and (ii) predict new combination therapies based on the differences in signaling networks between treated MS patients and controls (**Figure 1**). To this end, we assembled a literature- and database-based Prior Knowledge signaling Network (PKN) (21), which includes the pathways involved in immune and MS signaling, as well as their crosstalk and known therapeutic targets. Based on their ability to model large networks with a low number of parameters, logic models have been used to unravel the network topology driving disease and response to therapy (22–25). Here, we established a Boolean logic model from the signaling network and trained it with measurements of kinase de/phosphorylation as a proxy of signal propagation upon perturbation with ligands and drugs in Peripheral Blood Mononuclear Cells (PBMCs) from MS patients and healthy controls. We determined disease- and therapy-specific logic models that characterize the signaling networks for approved treatments (interferon-beta (IFNβ), glatiramer acetate (GA), natalizumab (NTZ), fingolimod (FTY) and the experimental drug epigallocatechin gallate (EGCG)). We hypothesized that the signaling interactions that the drugs failed to revert in the *ex vivo* assays to a healthy-like activity level may be candidates to be targeted by a second drug and hence lead to a personalized combination therapy. To identify those interactions, we developed a score of co-druggability of signaling interactions according to quantitative differences in network topology among healthy controls and untreated and treated MS patients (**Figure 1** and **Table 1**).

**Table 1.** Illustration of co-druggability score for extreme combinations of Healthy-MS-Treated signaling activity. Implications for each score using as an illustration extreme signaling activities. Based on the signaling activity of each interaction as assessed by modeling, the difference between healthy, untreated MS and treated MS (columns 1-3 for a given interaction) can be calculated, yielding a score (column 4). The implications for each score are shown in columns 5 and 6. Interactions with a negative co-druggability score indicated a treatment effect that produced signaling activity different to that found in healthy patients and were selected as co-druggable. A co-druggability score of 0 indicated that there was no effect due to drug treatment. From those cases, the interactions, in which signaling activity was different between healthy donors and treated patients were also selected as co-druggable, i.e. those where the drug alone failed to revert signaling to a healthy state. Hence, co-druggable interactions were defined as those where treatment with the drug alone yielded signaling activity different to that of the healthy-like state.

| Healthy | MS | Treated | Score | Treatment affecting? | Healthy-like signaling after treatment? |
| --- | --- | --- | --- | --- | --- |
| 0 | 0 | 0 | 0 | No | Yes |
| 0 | 0 | 1 | -1 | Yes | No |
| 0 | 1 | 0 | 1 | Yes | Yes |
| 0 | 1 | 1 | 0 | No | No |
| 1 | 0 | 0 | 0 | No | No |
| 1 | 0 | 1 | 1 | Yes | Yes |
| 1 | 1 | 0 | -1 | Yes | No |
| 1 | 1 | 1 | 0 | No | Yes |

Yes No
Effective Treatment Co-druggable (Treatment is affecting) Co-druggable (Treatment is not affecting)

**Figure 1.**
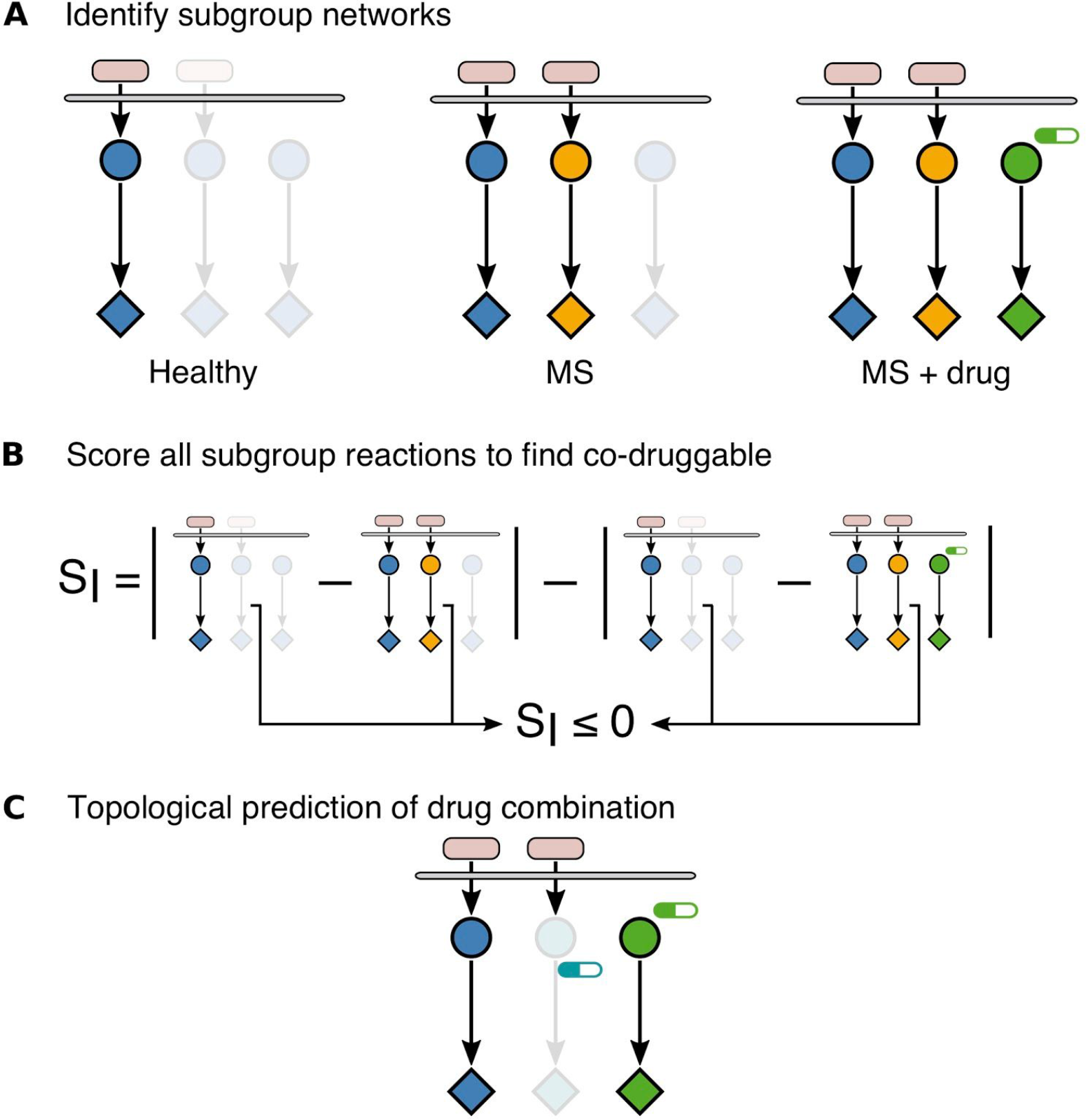
Topological modeling approach of signaling pathways for prediction of combination therapy. A) Identification of subgroup networks. A model characterizing signaling activity (upstream kinase (circle) that regulates a response e.g. innate immunity, survival (diamond)) in response to a stimulus (oval) was calculated for each donor based on the experimentally acquired dataset. Next, the donor-specific models are merged for all donors belonging to the same subgroup (left panel blue: healthy controls; middle panel orange: untreated MS; right panel green: treated MS). B) Scoring subgroup interactions to find co-druggable network interactions. The score is calculated to identify interactions that differ from healthy-like signaling activity in spite of drug treatment (see methods). C) Topological prediction of drug combination. A topology-based graph search allows identifying secondary treatments that could target and revert signaling of co-druggable interactions to a healthy-like activity state.

Using this network-based approach, we predicted several combination therapies, in particular of TAK1 with all five studied MS drugs. We validated a highly-scoring combination therapy in the animal model of MS experimental autoimmune encephalomyelitis (EAE). The modeling approach shown here can be used for designing combination therapies for other complex diseases as well as for developing personalized therapies.

## Results

### Multiplexed phosphoproteomic analyses in PBMCs from MS patients and controls

To characterize the signaling networks involved in MS, we created a Prior Knowledge Network (PKN) of biochemical interactions (kinase phosphorylation) reported to be involved in immune signaling associated with MS based on omics and functional studies, as well as those pathway interactions targeted by MS drugs (17). Our network includes pathways such as interferon response, B- and T- cell receptor signaling, cellular survival and apoptosis, lipid signaling, innate immunity and multi-drug response (MDR) genes (**Supplementary Figure S1**). To achieve this, we searched in state-of-the-art databases for interactions that were reported in highly specific assays and prioritized experiments with human and PBMC cells (**Supplementary Table S1 and S2**). Further, targets of MS drugs were included into the network via their crosstalk with immune pathways. The PKN featured 167 signaling components (**Supplementary Table S2**) and 294 interactions (**Supplementary Table S1**).

Subsequently, we developed a multiplex xMAP phosphoprotein panel tailored to logic modeling and aiming to maximize the coverage of the MS-specific network. The panel, which combined previously and newly developed assays, was then optimized to maximize accuracy, reproducibility as well as network coverage (see **Supplementary Table S3** and **Supplementary Methods S1**) (26). After optimization, we selected a set of 17 phosphoproteins with adequate signal to noise ratio that were used for the *in vitro* assays: AKT1, CREB1, FAK1, GSK3A, HSPB1, IKBA, JUN, MK12, MK03, MP2K1, PTN11, STAT1, STAT3, STAT5, STAT6, TF65, WNK1 (**Supplementary Table S4**). PBMCs were cultured in presence of different sets of stimuli such as lectins, endotoxins, immunostimulants, cytokines and drugs (**Supplementary Table S5**).

We extracted PBMCs from 195 MS patients (mean age: 43.1±11.3 years; disease duration: 8.7±7.7 years; median EDSS: 2.0 (0-6.0); subtype: 24 CIS, 129 RRMS, 6 SPMS and 36 PPMS; untreated: 98) and 60 healthy controls (**Supplementary Table S6**). From those 255 donors, 180 were selected and PBMCs were analyzed before stimulation and at 5 and 25 min after stimulation. This yielded a dataset consisting of three measurements for 17 phosphoproteins upon 20 stimuli, to a total of 183600 experimental measurements. For subsequent modeling, the two time points after baseline (5 and 25 min) were collapsed into a single activation signature to define whether a given phosphoprotein was activated or inhibited (see details in **methods**). Finally, we applied stringent quality control analyses of the dataset consisting of positive and negative controls, reproducibility tests as well as tests for artifacts in the distribution of the samples (see **Supplementary Methods S1**), which led to the removal of 2028 data points and 11 patients to generate the final dataset for a total of 169 subjects. After application of identifiability analysis based on the experimental coverage of our measurements (21), the PKN (shown in **Supplementary Figure S1**) was reduced to 71 proteins and immune modulators and 168 interactions.

### Logic modeling enables characterization of signaling networks in healthy donors and MS patients

Next, we sought to train the generic PKN to these phosphoproteomic measurements in a donor-specific manner to identify each donor’s active pathways. To train the same Boolean logic model to the data of each of the 169 subjects, the phosphoproteomic datasets were transformed (**Figure 2A**). To enable comparison of the data to the binary output of the Boolean model, we established a stringent normalization pipeline combining (i) a non-linear transformation to normalize the measurements to continuous values between 0 and 1 while penalizing the outliers (21) with (ii) a statistical filter that removed data-points belonging to proteins that were not found to be significantly phosphorylated or dephosphorylated (**Supplementary Figure S2**). **Figure 2B** shows the log fold change of each phospho-profile with respect to the control, **Figure 2C** the dataset after the non-linear normalization, and **Figure 2D** displays the proteins found to be significantly dephosphorylated or phosphorylated. Using this normalized dataset, we aimed to identify the active specific signaling network for each donor.

**Figure 2.**
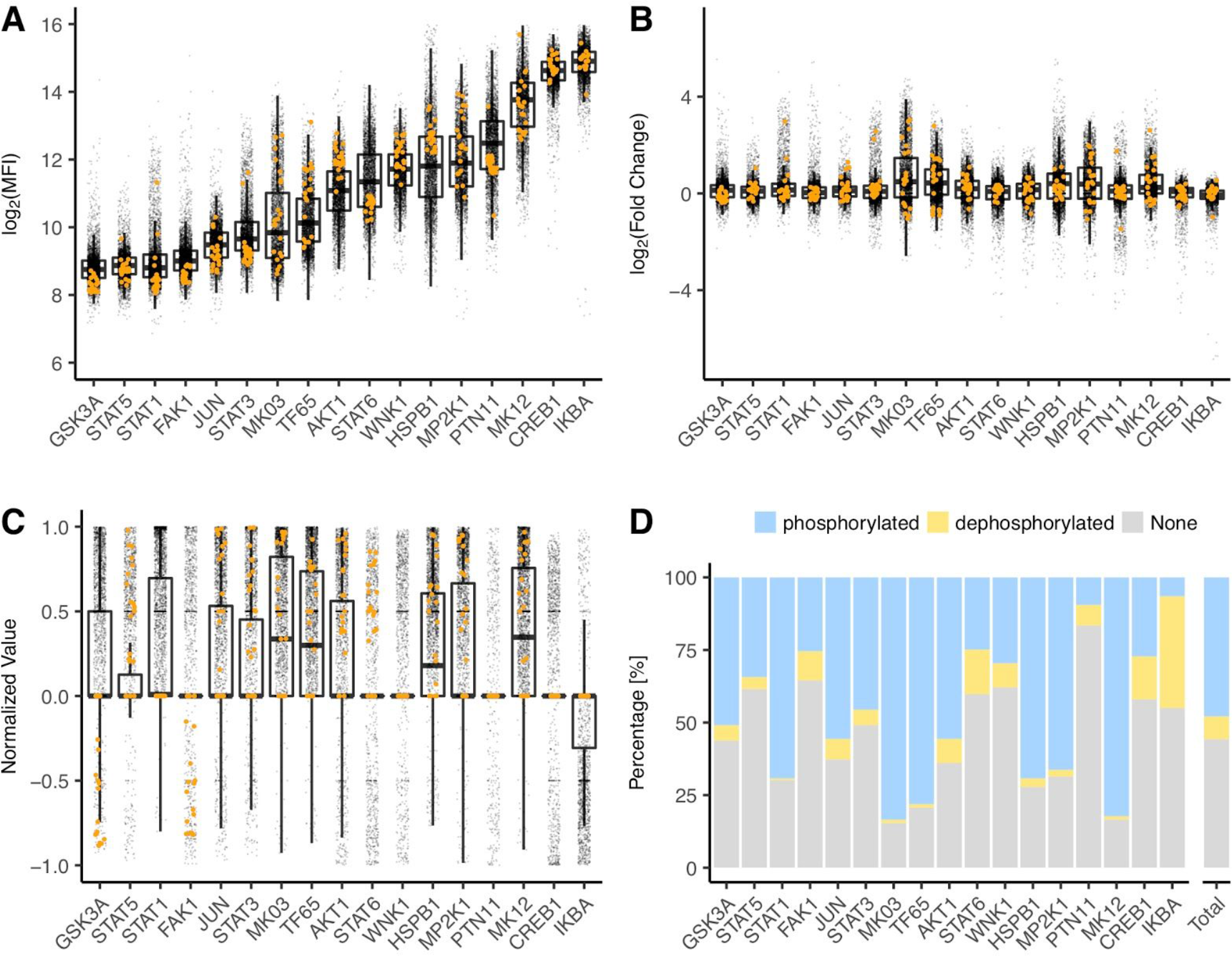
Phosphoproteomic measurement and normalization pipeline. **A)** xMAP Mean Fluorescence Intensity (MFI) log values of the 17 analyzed phosphoproteins; **B)** fold change distribution; **C)** non-linearly normalized values (see methods). Orange measurements A-C: values of the same patient to allow visualization of the changes across data transformation; **D)** Percentage of patients, for which each phosphoprotein was classified as phosphorylated, dephosphorylated or non-significant after statistical testing.

To this end, we fitted a general logic model derived from the PKN (21) to each donor dataset, which resulted in a compendium of 169 models. Model optimization was performed with CellNOpt, a tool that selects the logic model that best matches the data while penalizing model size (27). To increase the robustness of each model, optimization was performed 10 times, and for each donor we selected the best solution as well as all models with a data-to-simulation mismatch within a given relative tolerance reflecting that, due to lack of identifiability and technical noise, different models are feasible (21) (see **methods** for details). We then built a final model for each patient, defined by the median value of each interaction across all solutions within the relative tolerance (or borders of tolerance) of the best solution (**Figure 3A**). To assess the validity and robustness of our modeling approach, we confirmed absence of bias due to treatment, center, disease subtype, medical condition, as well as technical aspects such as model size and merging strategy (**Supplementary Figure S3**).

**Figure 3.**
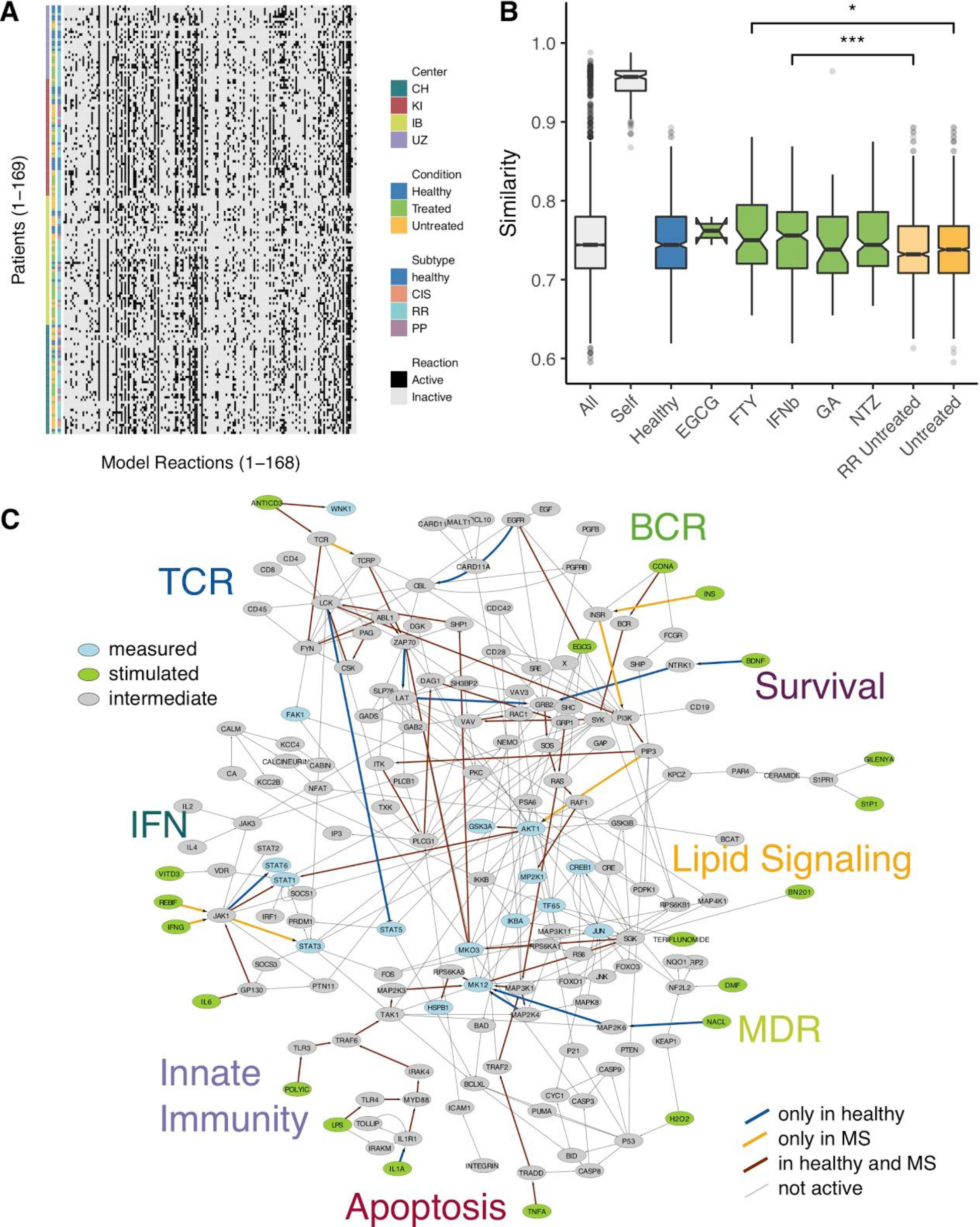
Logic modeling identifies donor-specific signaling networks and reveals MS-specific signaling pathways. **A)** Signaling network found by modeling for each donor, visualized as a heatmap. Rows: Single donor network. Columns: Signaling activity determined for each interaction by calibrating the PKN shown in Supplementary Figure S1 after removing the unidentifiable interactions using the phosphoproteomics dataset of each donor. **B)** Simple Matching Coefficient (SMC) was used to quantify background similarity among all models (All), each model with all solutions for a single donor (Self), and within donor subgroups (MS drugs: FTY, IFNβ, GA, NTZ; experimental drug: EGCG, untreated RRMS (for direct comparison with MS drugs subgroups) and untreated MS (for comparison with healthy controls)). Similarity is higher within each subgroup than compared to background. **C)** Differentially activated pathways (see Supplementary Methods) between healthy controls (HC) and untreated MS patients (MS). The models previously calculated for each donor were merged to reveal the common active pathways for controls (blue), untreated MS patients (orange) and both (brown). Grey: Inactive interactions from the MS, immune- and treatment-related network (Supplementary Figure S1).

To confirm that our modeling approach was able to capture the effect on signal transduction of MS drugs, intra-as well as inter-donor model similarities were calculated by the Simple Matching Coefficient (SMC) index. This means, we assessed the number of interactions in two networks that are equal in both networks over the number of all interactions, after stratification of patients by treatment subgroups (**Figure 3B**). Modeling shows that treated MS patients (green boxes) yield more similar signaling activity than untreated MS patients (orange boxes). Treatment with FTY yielded the overall highest within-subgroup similarity and was statistically more similar in this group’s patients than compared to RRMS untreated patients (untreated RRMS mean similarity: 0.737, FTY mean similarity: 0.755, Wilcoxon’s test p-value 0.0025). Within-subgroup similarity was also higher for patients treated with IFNβ (untreated RRMS mean similarity: 0.737, IFNβ mean similarity: 0.750, Wilcoxon’s test p-value 7.81*10^−6^). The large similarity of all solutions found during model optimization for each patient (mean similarity: 0.949) further supported the use of an individual model for each donor. Altogether, these results supported that merging models by subgroups of donors would yield biologically meaningful signaling networks for each group.

We then merged the individual models of healthy donors and separately those of untreated MS patients to characterize MS signaling. We calculated the mean of each interaction for each subgroup and removed the inactive interactions, i.e. those with an activity value below the upper quartile, which we considered neglectable. The differentially activated interactions between healthy donors and untreated MS patients uncovered signaling pathways deregulated in MS (**Supplementary methods** and **Figure 3C)**. Our analysis revealed the activation of several pathways in PBMCs after stimulation both in patients and controls, ranging from cell survival and proliferation to TCR, innate immunity, and pro-inflammatory response pathways (e.g. TCR - CSK - LCK, JAK1 - STAT1, TLR4/IL1R1 - MYD88 - IRAK4 - TRAF6 - TAK1, TNFα - TRADD - TRAF2 - MAP3K1, SGK - MK03, or RS6 - SGK, RAF1 - MP2K1). Further, pathways such as INSR - PI3K - PIP3 - AKT1 and JAK1 - STAT3 were found to be activated in MS patients, while BDNF - NTRK1 - GRB2 was found to be activated in healthy donors.

These findings indicated that our method was able to identify previously described pathways in the setting of *ex vivo* analysis of human PBMCs (17), as examined in detail for the individual pathways in the discussion. Untreated MS patients, when compared to healthy controls, showed an increase in the activation of the NFkβ pathway (TAK1-IKKB), the activation of the cell prosurvival PI3K pathway (SLP76-AKT1) or the activation of interferon/cytokines pathways (JAK1-STAT3). Moreover, patients showed a decrease, compared to healthy controls, in the activation of TCR/IL-2 pathway (LCK-STAT1) as well as in the effect of trophic factors signaling on ubiquitination system (EGFR-CBL). This results characterize at the mechanistic and quantitative level the differential activation of the immune pathways in MS.

### Quantitative differences found in signaling pathways under therapy

We then quantified differences in signaling between patients treated with MS-specific therapies, using the same procedure as for healthy donors versus untreated MS patients. We focused on the strongest signals, i.e. those in the upper quartile of the group mean (**Supplementary Figure S4**). This yielded a signaling network of active reactions characterizing each subgroup, uncovering the effect of each MS-treatment on signaling at the mechanistic level (**Supplementary Figure S5**).

Interestingly, the activation of TAK1, a key component involved in TGF-β, toll-like receptor, B-cell receptor and NFkβ pathways, as well as response to inflammation was identified in all 5 networks inferred under treatment (**Supplementary Figure S5A-E**). Additionally, its activity was persistently over-activated in terms of quantitative signaling activity by the currently approved MS drugs (**Supplementary Figure S4**). In the case of IFNβ and GA treated patients, the network revealed the activation of STAT1 and STAT3 by JAK1, whereas in the EGCG, FTY and NTZ specific networks the activation of the JAK1-STAT pathway is regulated via STAT3, STAT6 and STAT1 respectively.

To validate that the therapy-specific pathways found were consistent with the experimental measurements of MS signatures, we quantified the degree to which the original phospho-measurements supported the signaling pathways predicted by modeling for each patient subgroup. The proteins differentially phosphorylated across all stimuli combinations were overrepresented in pathways found by model fitting (IFNβ p=1.2E-05; GA p=1.4E-06; FTY p=0.0004; NTZ p=0.006; Fisher’s exact test) except for EGCG, which was limited by the small sample size (**Supplementary Figure S5F** and **Methods**).

### Network topology-based prediction of targeted combination therapy

Our main goal was to use the subgroup networks found to predict novel combination therapies. To this aim, we defined as therapeutic goal to revert the signaling network of patients with MS to a healthy-like activity. We introduced a restriction to our method: the combinations we sought featured one approved MS drug which, in spite of its known efficacy in MS, yielded a signaling activity that differs from the healthy controls as identified by our topological modeling approach. Thereby, we aimed to predict which kinase interactions within the network would revert signaling activity to the healthy-like state when combined with the ongoing therapy. We hypothesized that a co-druggable interaction, i.e. those that a given MS therapy failed to revert to a healthy-like activity level, should be the model interactions with a signaling value more distant between healthy control and MS drug models than between healthy control and untreated MS models (**Figure 1** and **Methods**). This yielded a list of interactions with their corresponding score quantifying co-druggability potential **(Supplementary Table S7** and **Supplementary Figure S4**). To identify the co-druggable interactions, we selected those with a non-positive score as described above, and filtered for additional conditions (see **Methods**), thereby identifying both the interactions whose signaling activity was different from healthy because of deregulation by the disease as well as because of off-target primary drug effect **(Figure 4A)**. The last step of our approach was to predict the stimulus that would revert to healthy-like signaling activity of those interactions identified as co-druggable. Therefore, the co-druggable interactions were mapped onto the signaling network assessed for each drug subgroup. As an example, **Figure 4B** shows the co-druggable interactions under FTY treatment mapped onto the FTY network. Mapping the co-druggability for each interaction allowed us to predict combination therapies based on each treatment’s signaling network topology. Finally, we employed a graph search approach to identify the *in vitro* stimulus used in our study that activated interactions found to be co-druggable with each drug. In other words, we found the experimental readouts that could be measured and reached from an *in vitro* stimulus via an interaction found to be co-druggable *in vivo* (**Supplementary Table S8**). It is important to note that our approach enabled the prediction of combinations of approved MS therapies which yielded a signaling activity distant from healthy controls with drugs that stimulated or inhibited signaling. Among all predictions (**Supplementary Table S8**), we chose to validate the combination of FTY with a TAK1 inhibitor based on (i) the striking signaling homogeneity across FTY models (as quantified in **Figure 3B**), (ii) the finding that TAK1 modulation of HSPB1 via MK12 is largely deregulated (**Supplementary Figure S4**, panel FTY, interaction MK12 - HSPB1) and (iii) the fact that TAK1 is active and co-druggable in all 5 networks under treatment with *in vivo* drugs, specifically TAK1 - MAP2K4 for ECGC, IL1R1 - TAK1 for FTY, IL1R1 - TAK1 for GA, LPS - TAK1 for IFNβ and TAK1 - MK12 for NTZ (**Supplementary Table S7**).

**Figure 4.**
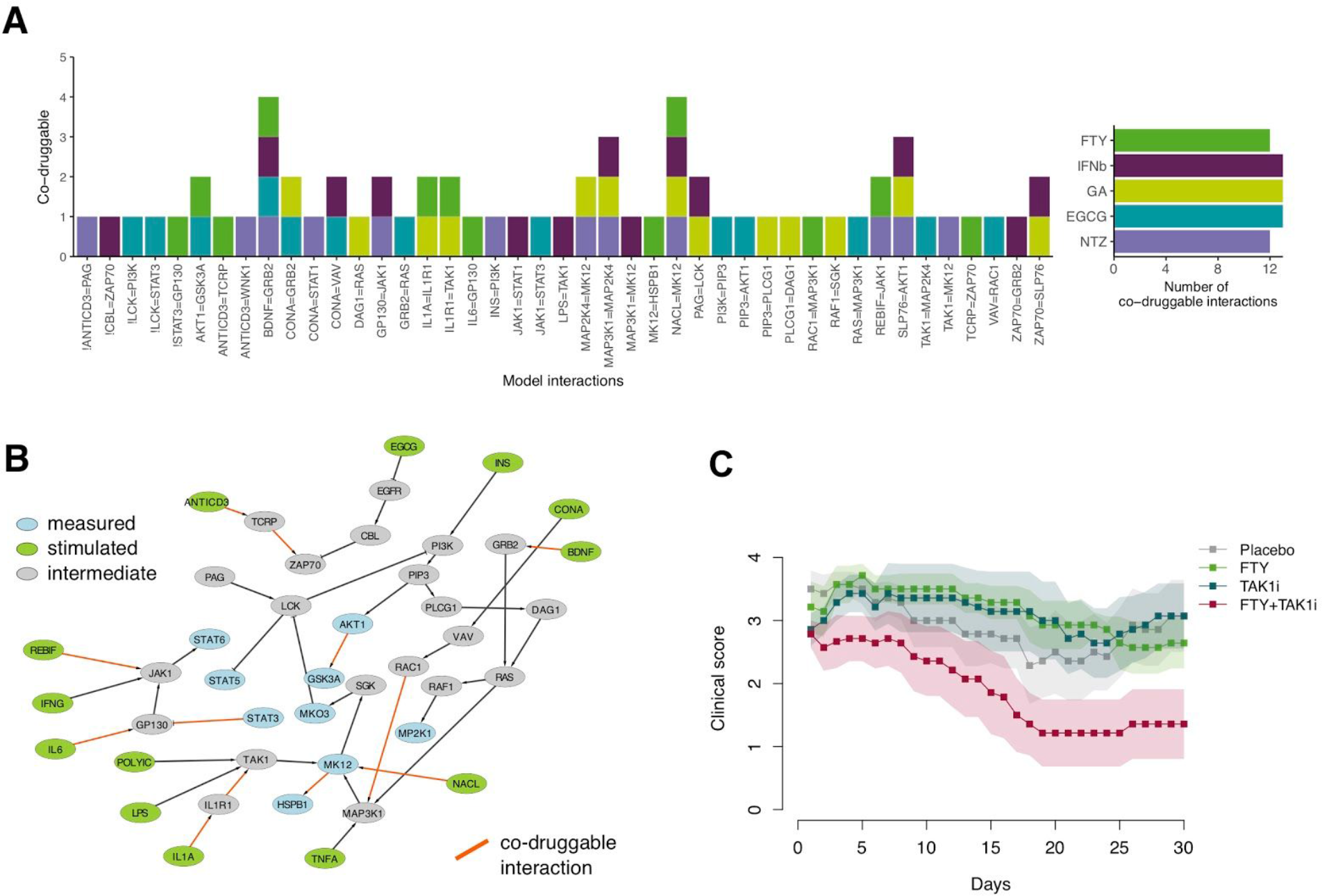
Combination therapies predicted and *in vivo* validation. **A)** All predicted co-druggable interactions of the MS drugs models. Based on the subgroup models, the co-druggability of all 168 network interactions (X axis) was assessed using the co-druggability score, and those identified as co-druggable (see Figure 1, Table 1 and main text) are shown. For each interaction (X axis) the number of drugs (Y axis) is shown, in which it was found to be co-druggable using the co-druggability criteria; **B)** FTY network co-druggability: the case of FTY network co-druggability is shown as an example (red line: interactions predicted to be co-druggable); **C)** *In vivo* validation of the combination FTY+TAK1-inhibitor in the EAE model. The graph shows the mean and the standard error of the clinical score for each group (n=10). Animals started treatment after disease onset (clinical score >1.0) and were randomized to each treatment and rated in a blinded manner.

### Validation of a predicted combination therapy in the animal model of MS

To validate our approach and obtain an *in vivo* proof-of-concept of the efficacy of the combination therapies proposed, we sought to assess whether the combination therapies improved the clinical course of the animal model of MS (EAE in the C57BL6 mice immunized with the MOG_35-55_ peptide). Mice were randomized after disease onset (after they reached a clinical score >1.0), either to placebo (saline), each drug alone (using doses previously demonstrated as efficacious in rodent models) or the combination of FTY with the TAK1 inhibitor (5Z-7-oxozeaenol). We found that the combination FTY+TAK1-inhibitor ameliorated the clinical course of EAE compared to placebo, TAK1-inhibitor or FTY alone (Mann-Whitney test; p<0.05) (**Figure 4C**). Therefore, our method was able to identify combinations of drugs targeting different pathways that achieved higher efficacy than single therapy.

## Discussion

In this study, we applied a modeling approach to characterize signaling activity both at the donor-specific and at the subgroup level using phosphoproteomic data from primary immune cells of MS patients and healthy donors. At the subgroup level, we identified models of the signaling network for healthy controls, untreated and drug-treated MS patients. Our approach allowed us to characterize signaling deregulation in this complex and heterogeneous disease. Further, based on the data-driven models, we developed a network-based method to predict novel combination therapies for the treatment of MS. We hypothesized that the interactions in the subgroup models in response to single drugs that were not reverted to a healthy-like activity state can be co-drugged. Hence, we developed a co-druggability score identifying those interactions. Finally, we used that score on a newly developed strategy to predict combination therapies based on network topology. Thus, our algorithm identifies interactions within signaling networks that should be targeted for restoring the network to a healthy-like state.

As validation, we tested the combination of one approved drug for MS, FTY, with an inhibitor of TAK1, a key component in the predicted signaling networks of all five studied *in vivo* drugs, which was found highly deregulated in FTY. This combination largely ameliorated the course of the animal model of MS with significantly higher efficacy than each treatment alone, providing *in vivo* proof-of-concept for our approach.

The approach presented here allows insights into signaling deregulation in MS by comparing signaling networks in untreated MS patients with those of healthy controls. The comparison demonstrates enhanced pro-survival effects of the trophic factor signaling pathway (AKT1) and modulation of the interferon pathway (JAK1, STAT3). In line with the differences found here, aberrant STAT phosphorylation signaling in peripheral blood mononuclear cells from MS patients has been reported (28). The NFkβ pathway has been reported to be overactivated in PBMCs from patients with MS (29, 30) as well as to contribute to the genetic susceptibility of the disease (31, 32). Our analysis supports the systemic pro-inflammatory state of PBMCs in MS. The trophic factor pathway involving SLP76 and AKT was found active in patients under treatment with GA, NTZ, and IFNβ and has also been associated with MS (17) and MS susceptibility via CD6 gene (33, 34). The involvement may reflect a pro-survival signaling state of T- and B-cells in the context of the pro-inflammatory microenvironment. Finally, we observed overactivation of the cytokine/interferon pathway (JAK1), which has previously been reported in MS (17). Moreover, STAT3 has been confirmed as susceptibility gene for the disease (35), and its activation is impaired in response to IL-10 in MS patients (36), suggesting a defective response of regulatory Tr1 cells. Regarding the interactions that were decreased in MS patients compared to controls, PBMCs from patients with MS showed lowered inhibition of STAT5 by LCK suggesting impairment of the regulation of T/B-cell signaling and IL-2 trophic effects (37) or cytotoxicity (38), as well as the regulation of the ubiquitination system modulated by CBL-B (39). LCK is modulated by EVI5, and influences STAT5 in our analysis, both being susceptibility genes for MS (35). The second inhibited interaction involved the MS susceptibility gene CBL-B (40), which regulates TCR and co-stimulatory signals and regulates immune tolerance through its ubiquitin E3-ligase activity. CBL-B expression is reduced in CD4 cells from MS patients and alters the signaling of the type I interferon pathway (41, 42). CBL-B is activated by EGFR, leading to inhibition of several pathways by ubiquitination, including that of EGFR itself (43, 44). In summary, our results are supported by multiple previous studies identifying deregulation of pathways in MS (17) and shows the activation of pro-inflammatory pathways and the inhibition of pathways related with immune tolerance.

Regarding the signaling models of MS drugs, our method can identify the activation of pathways downstream of drug targets in our PKN. Further, it revealed interactions with other disease-associated pathways, which may not directly be involved in the signaling elicited by such drug. Therefore, our modeling approach can be used to provide a wider context of the effects of a given drug on the functions of the target cells. For example, the models identified the activation of JAK1-STAT1 pathway (45, 46) in IFNβ-treated patients, the activation of AKT, PLC (47, 48) in patients treated with FTY, or the activation of MAPK, NFkβ (via TAK1) (49–51) in NTZ treated patients.

Our study has several limitations. First, the analysis of signaling signatures based on phosphoproteomics of mixed immune cells can give rise to high signal variability that is confounded by the presence of multiple cell types. The analysis of signaling abnormalities from specific immune cell subtypes e.g. CD4+, CD8+, B-cells, monocytes that probably differentially contribute to the disease susceptibility or to the response to therapy may reveal new therapeutic targets. This limitation can be overcome by technologies that allow to collect information at the single-cell level such as mass cytometry in future studies (52), although the cost is considerably higher, thus limiting number of subjects. Second, our current coverage using validated xMAP assays was restricted to less than 20 kinases, a small subset of signaling molecules and pathways that may be activated in immune cells. Robust quantitative phosphoproteomic assays covering thousands of phosphosites in hundreds of samples can be used for logic modeling (53). This will allow us to expand significantly the scope of our models. Third, the Boolean logic approach does not describe processes biochemically and models processes as binary, hence missing subtle aspects of signal transduction (10). An important goal of our studies was to develop a modeling approach that can guide the rational development of combination therapies. Our signaling topology-based approach has the advantage of allowing prediction of combinations between drugs currently used in patients with MS and compounds that can stimulate signaling where those drugs yield a signaling activity distant from the healthy state. Many strategies have been developed to predict drug combinations (54). Others have used phosphorylation data upon perturbation using statistical approaches (13), data-driven network inference (5, 11), or a combination of mechanistic and Bayesian network modeling (55). Due to their simplicity, logic networks can model large networks and provide a useful framework to study drug combinations (23–25). We used this logic framework to analyze a large dataset encompassing 183600 data points derived from a newly recruited cohort of 169 donors. Our approach is a compromise between data availability, technical feasibility and computational burden that sacrifices details to be able to capture a broad portion of the signaling machinery. Fourth, in this study, we have explored only some of the currently approved drugs for MS patients. And fifth, we have averaged individual patients’ networks to obtain the subgroup (drug) networks, which leads to losing individual variability, probably related to differential genetic susceptibility or immune system activation.

In summary, we have built donor-specific dynamic logic models and developed a network-based approach to (i) characterize MS and treatment deregulation of signaling and (ii) predict new targets for combination therapy. This approach can be applied to other diseases.

## Materials and Methods

### Subjects and clinical cohorts

We recruited 255 subjects including 195 patients with MS and 60 healthy controls, matched for age and sex with the RRMS group, in a multi-centric study in four MS centers (Hospital Clinic Barcelona – IDIBAPS (n=69), Karolinska Institute (n=64), University of Zurich (n=40) and Charité University (n=82)) (**Supplementary Table S5** and **Supplementary Methods S1**).

### Samples and processing

A unified standard operating procedure for PBMCs isolation, stimulation and lysis, as well as sample storing and shipping was developed along with a kit (plates) with reagents and buffers that were produced in a single facility (ProtAtOnce) and shipped to all participating centers (see **Supplementary Methods S1** for details). The reagents were prepared from a single batch and plates were prepared from a single batch for each stimulus. Quality controls were carried out to ensure that the reagents remain stable for 3 months.

### xMAP assays

XMAP assays were developed by ProtAtOnce (Athens, Greece) and were standardized to minimize error. We optimized assays from a list of 70 candidates (see **Supplementary Methods S1**) and obtained a final list of 17 phosphoproteins which display a good signal to noise ratio to be measured in the *in vitro* assays: AKT1, CREB1, FAK1, GSK3A, HSPB1, IKBA, JUN, MK03, MK12, MP2K1, PTN11, STAT1, STAT3, STAT5A, STAT6, TF65, WNK1 (**Supplementary Table S3** and **S4**). We used a set of 20 stimuli, which included pro-inflammatory or pro-oxidant stimuli (Anti-CD3, concanavaline A (conA), H2O2, IFNG, IL1A, IL6, LPS, NaCl, PolyIC, TNFα), immunomodulatory stimuli (S1P, vitD3) neuroprotectants or anti-oxidants (BDNF, EGCG, INS and BN201), disease modifying drugs from MS (DMF, FTY, Teriflunomide, IFNβ1a (Rebif®)), and a culture media as control (**Supplementary Table S5**). Samples were collected at baseline (time 0) and after 5 and 25 min.

### Data normalization

After signal reading, data was normalized. To allow logic modeling, data was normalized between 0 and 1, extending via stringent statistics the normalization strategy presented in (21) (see **Supplementary Methods S1**). The maximum between 5 and 25 measurements was selected to allow capturing signal transduction including late effects of network motifs such as negative feedback loops.

### Model generation

Healthy controls and MS (untreated) signaling models as well as MS drug-specific models were assembled, using manual curation to prioritize interactions that were found in immune or human cells whenever possible. To allow inclusion of MS drugs, the drug targets were included in pathways as explained above via the references detailed in **Supplementary Tables S1** and **S2**. The global network featured (167 nodes and 294 interactions). The identifiable components, i.e. those that led to unique solutions using the optimization algorithm, were assessed as described in (21), yielding a so-called preprocessed network of 71 nodes and 168 interactions. To reduce model complexity to a degree that could be solved by model optimization, AND gates were added based on publicly available knowledge on protein complexes (see **Supplementary Methods S1**).

### Model Optimization

For each patient, 10 completed optimization runs were performed with a genetic algorithm, and assessed as shown in **Supplementary Figure S3** (see **Supplementary Methods S1**).

### Network topology-based prediction of combination therapy

We calculated the co-druggability score as described in **Figure 1** with *S* indicating the signaling activity status of each interaction:

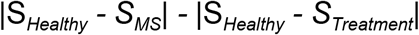

were *Healthy* refers to the healthy subgroup model, *MS* relates to the untreated MS subgroup model and *Treatment* denotes one of the five approved drug treated MS subgroup models. Therefore, interactions with a negative co-druggability score indicated a treatment effect that produced signaling activity more different from that found in healthy donors than without treatment and were selected as co-druggable. A co-druggability score of 0 indicated that there was no effect due to drug treatment.

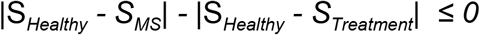

Thus, we used the algorithm described in **Figure 1** to define the co-druggability of all interactions within each treatment group after filtering for active signaling. To ensure that interactions with positive scores close to zero were also captured by our method (interactions with a similar signaling activity between drug treatment and MS untreated), we defined a lower quartile threshold around zero and collapsed almost zero scores into zero. Further, we required co-druggable interactions to be significantly (p<0.05) different between healthy and drug signaling by using the upper quartile as difference threshold. Applying both filters together allowed us to identify co-druggable interactions that were deregulated by the disease as well as the unwanted signaling effect by the primary drug (**Figure 4** and **Supplementary Figure S4**).

### Experimental Autoimmune Encephalomyelitis

Female C57BL/6 mice from Harlan (8-12 weeks old) were immunized subcutaneously in both hind pads with 300 µg of a myelin oligodendrocyte glycoprotein (MOG_35-55_, Spikem, Florence, Italy) emulsified with 50 µg of *Mycobacterium tuberculosis* (H37Ra strain; Difco, Detroit, MI, US) in incomplete Freund’s adjuvant as described previously (56). Mice were injected intraperitoneally with *Pertussis toxin* (500 ng; Sigma, US) at the time of immunization and 2 days later. Animals were weighed and inspected for clinical signs of disease on a daily basis by an observer blind to the treatments. The severity of EAE was assessed on the following scale: 0= normal; 0.5= mild limp tail; 1= limp tail; 2= mild paraparesis of the hind limbs, unsteady gait; 3= moderate paraparesis, voluntary movements still possible; 4= paraplegia or tetraparesis; 5= moribund state; 6= death. Animals (10 mice per arm) were randomized to their treatment once they have reached a clinical score > 1 point. Clinical assessment was performed by a blinded evaluator to the treatment group. Comparison between groups were performed using the Mann-Whitney test with p-value cut-off at 0.05.

### Statistical validation of patient subgroup models

To test whether the signaling models found for each patient subgroup were reflected in the experimental phospho-levels, we first calculated for each stimuli pair, if the fold change values (before non-linear normalization) within a single subgroup deviated significantly from 0 by a one-sample Wilcoxon test. Multiple testing correction was performed using the Benjamini-Hochberg strategy. Secondly, we grouped the stimulus pairs into two categories, i.e. either in accordance with the model of signaling or not, by confirming if a phospho-readout could be reached by the respective stimulus in the signaling model for the different subgroups (**Supplementary Figure S5A-E**). Finally, using Fisher’s exact test, we then assessed if the stimuli pairs found supported by the models were enriched for high significance (using p<0.05 as cutoff) compared to the not supported pairs (**Supplementary Figure S5F**).

### Code and data availability

All code, mathematical models and data can be found on Github (http://github.com/saezlab/combiMS).

## Acknowledgements

The authors thank Federica Eduati for insightful discussion and valuable advice all along the way. We also would like to thank Thomas Cokelaer for crucial technical help. Funding: This work was supported by the European Union 7FP-Programme (CombiMS, grant No 305397) and the European Sys4MS project (Horizon2020: Eracosysmed: ID-43). RM and the Section for Neuroimmunology and MS Research, University Zurich (UZH), have been supported by a European Research Council Advanced Grant (ERC 340733-HLA-DR15 in MS), by a Clinical Research Priority Project - disease heterogeneity of MS (CRPP^MS^; UZH), by the Swiss National Science Foundation and the Swiss MS Society.

## Author contributions

Developed the modeling and combination therapy prediction approach: MBF. Conceived and designed the experiments: MBF, IP, VP, DEM, LGA, JSR, PV. Developed new phospho-assays: VP, DEM, TS, LGA. Designed patient cohort and recruited patients: TO, RM, FP, PV. Performed the experiments: VP, DEM, GV, WF, PS, JB. Contributed reagents/materials/analysis tools: MBF, TO, RM, FP, LGA, JSR, PV. Analyzed the data: MBF, JW, TS, JSR. Tested algorithm, performed statistical validation and additional analyses and modified figures accordingly: JW, MR. Wrote the manuscript: MBF. Edited the manuscript: JSR, PV. Supervised the clinical study and experimental validation: PV. Supervised the modeling, combination therapy prediction and overall project: JSR. All authors revised and approved the final manuscript.

## Competing interest

DEM is an employee of ProtATonce; TO has received honoraria for lectures and/or advisory boards as well as unrestricted Multiple Sclerosis research grants from Allmiral, Astrazeneca, Biogen, Genzyme, Merck and Novartis; RM reports grants and personal compensation from Biogen, personal fees from Genzyme Sanofi Aventis, grants and personal fees from Novartis, personal fees from Merck Serono, Roche, Neuway and CellProtect, outside the submitted work; FP has received research grants and personal compensation for activities with Alexion, Bayer, Chugai, Novartis, Merck, Teva, Sanofi, Genzyme, Biogen and MedImmune; LGA is founder and hold stocks at ProtATonce; PV holds stocks and has received consultancy fees from Bionure Farma SL, Spiral Therapeutics Inc, Spire Bioventures Inc, Attune Neurosciences Inc, QMenta Inc and Health Engineering SL; All other authors have nothing to disclose.

## References

1. Owens J (2007) Determining druggability. Nat Rev Drug Discov 6(3):187–187.

2. Jia J, et al. (2009) Mechanisms of drug combinations: interaction and network perspectives. Nat Rev Drug Discov 8(2):111–128.

3. Cully M (2015) Combinations with checkpoint inhibitors at wavefront of cancer immunotherapy. Nat Rev Drug Discov 14(6):374–375.

4. Klinger B, et al. (2013) Network quantification of EGFR signaling unveils potential for targeted combination therapy. Mol Syst Biol 9:673.

5. Bozic I, et al. (2013) Evolutionary dynamics of cancer in response to targeted combination therapy. Elife 2:e00747.

6. Villoslada P, Steinman L, Baranzini SE (2009) Systems biology and its application to the understanding of neurological diseases. Ann Neurol 65(2):124–139.

7. Vaga S, et al. (2014) Phosphoproteomic analyses reveal novel cross-modulation mechanisms between two signaling pathways in yeast. Mol Syst Biol 10(12):767–767.

8. Kholodenko BN, Hancock JF, Kolch W (2010) Signalling ballet in space and time. Nat Rev Mol Cell Biol 11(6):414–426.

9. Kolch W, Halasz M, Granovskaya M, Kholodenko BN (2015) The dynamic control of signal transduction networks in cancer cells. Nat Rev Cancer 15(9):515–527.

10. Davis MM, Tato CM, Furman D (2017) Systems immunology: just getting started. Nat Immunol 18(7):725–732.

11. Korkut A, et al. (2015) Perturbation biology nominates upstream-downstream drug combinations in RAF inhibitor resistant melanoma cells. Elife 4. doi:10.7554/eLife.04640.

12. Palacios R, et al. (2007) A network analysis of the human T-cell activation gene network identifies JAGGED1 as a therapeutic target for autoimmune diseases. PLoS One 2(11):e1222.

13. Lee MJ, et al. (2012) Sequential application of anticancer drugs enhances cell death by rewiring apoptotic signaling networks. Cell 149(4):780–794.

14. Ransohoff RM, Hafler DA, Lucchinetti CF (2015) Multiple sclerosis-a quiet revolution. Nat Rev Neurol 11(3):134–142.

15. International Multiple Sclerosis Genetics Consortium, et al. (2011) Genetic risk and a primary role for cell-mediated immune mechanisms in multiple sclerosis. Nature 476(7359):214–219.

16. Bayat Mokhtari R, et al. (2017) Combination therapy in combating cancer. Oncotarget 8(23):38022–38043.

17. Kotelnikova E, et al. (2015) Signaling networks in MS: a systems-based approach to developing new pharmacological therapies. Mult Scler 21(2):138–146.

18. Kieseier BC, Stüve O (2011) Multiple sclerosis: combination therapy in MS--still a valid strategy. Nat Rev Neurol 7(12):659–660.

19. Conway D, Cohen JA (2010) Combination therapy in multiple sclerosis. Lancet Neurol 9(3):299–308.

20. Milo R, Panitch H (2011) Combination therapy in multiple sclerosis. J Neuroimmunol 231(1-2):23–31.

21. Saez-Rodriguez J, et al. (2009) Discrete logic modelling as a means to link protein signalling networks with functional analysis of mammalian signal transduction. Mol Syst Biol 5:331.

22. Bernardo-Faura M, Massen S, Falk CS, Brady NR, Eils R (2014) Data-derived modeling characterizes plasticity of MAPK signaling in melanoma. PLoS Comput Biol 10(9):e1003795.

23. Eduati F, et al. (2017) Drug Resistance Mechanisms in Colorectal Cancer Dissected with Cell Type-Specific Dynamic Logic Models. Cancer Res 77(12):3364–3375.

24. Silverbush D, et al. (2017) Cell-Specific Computational Modeling of the PIM Pathway in Acute Myeloid Leukemia. Cancer Res 77(4):827–838.

25. Flobak Å, et al. (2015) Discovery of Drug Synergies in Gastric Cancer Cells Predicted by Logical Modeling. PLoS Comput Biol 11(8):e1004426.

26. Poussin C, et al. (2014) The species translation challenge-a systems biology perspective on human and rat bronchial epithelial cells. Sci Data 1:140009.

27. Terfve C, et al. (2012) CellNOptR: a flexible toolkit to train protein signaling networks to data using multiple logic formalisms. BMC Syst Biol 6:133.

28. Canto E, et al. (2018) Aberrant STAT phosphorylation signaling in peripheral blood mononuclear cells from multiple sclerosis patients. J Neuroinflammation 15(1):72.

29. Moreno B, et al. (2006) Methylthioadenosine reverses brain autoimmune disease. Ann Neurol 60(3):323–334.

30. Chen D, et al. (2016) CD40-Mediated NF-κB Activation in B Cells Is Increased in Multiple Sclerosis and Modulated by Therapeutics. J Immunol 197(11):4257–4265.

31. Housley WJ, et al. (2015) Genetic variants associated with autoimmunity drive NFκB signaling and responses to inflammatory stimuli. Sci Transl Med 7(291):291ra93.

32. Hussman JP, et al. (2016) GWAS analysis implicates NF-κB-mediated induction of inflammatory T cells in multiple sclerosis. Genes Immun 17(5):305–312.

33. Johnson BA, et al. (2010) Multiple sclerosis susceptibility alleles in African Americans. Genes Immun 11(4):343–350.

34. Hassan NJ, et al. (2006) CD6 regulates T-cell responses through activation-dependent recruitment of the positive regulator SLP-76. Mol Cell Biol 26(17):6727–6738.

35. International Multiple Sclerosis Genetics Consortium (IMSGC), et al. (2013) Analysis of immune-related loci identifies 48 new susceptibility variants for multiple sclerosis. Nat Genet 45(11):1353–1360.

36. Martinez-Forero I, et al. (2008) IL-10 suppressor activity and ex vivo Tr1 cell function are impaired in multiple sclerosis. Eur J Immunol 38(2):576–586.

37. Beyer T, et al. (2011) Integrating signals from the T-cell receptor and the interleukin-2 receptor. PLoS Comput Biol 7(8):e1002121.

38. Slavin-Chiorini DC, et al. (2004) Amplification of the lytic potential of effector/memory CD8+ cells by vector-based enhancement of ICAM-1 (CD54) in target cells: implications for intratumoral vaccine therapy. Cancer Gene Ther 11(10):665–680.

39. Swaminathan G, Tsygankov AY (2006) The Cbl family proteins: ring leaders in regulation of cell signaling. J Cell Physiol 209(1):21–43.

40. Sanna S, et al. (2010) Variants within the immunoregulatory CBLB gene are associated with multiple sclerosis. Nat Genet 42(6):495–497.

41. Stürner KH, Borgmeyer U, Schulze C, Pless O, Martin R (2014) A Multiple Sclerosis–Associated Variant of CBLB Links Genetic Risk with Type I IFN Function. The Journal of Immunology 193(9):4439–4447.

42. Sellebjerg F, Krakauer M, Khademi M, Olsson T, Sørensen PS (2012) FOXP3, CBLB and ITCH gene expression and cytotoxic T lymphocyte antigen 4 expression on CD4(+) CD25(high) T cells in multiple sclerosis. Clin Exp Immunol 170(2):149–155.

43. Galisteo ML, Dikic I, Batzer AG, Langdon WY, Schlessinger J (1995) Tyrosine phosphorylation of the c-cbl proto-oncogene protein product and association with epidermal growth factor (EGF) receptor upon EGF stimulation. J Biol Chem 270(35):20242–20245.

44. Tarcic G, et al. (2009) An unbiased screen identifies DEP-1 tumor suppressor as a phosphatase controlling EGFR endocytosis. Curr Biol 19(21):1788–1798.

45. Pertsovskaya I, Abad E, Domedel-Puig N, Garcia-Ojalvo J, Villoslada P (2013) Transient oscillatory dynamics of interferon beta signaling in macrophages. BMC Syst Biol 7:59.

46. Schneider WM, Chevillotte MD, Rice CM (2014) Interferon-stimulated genes: a complex web of host defenses. Annu Rev Immunol 32:513–545.

47. Takuwa Y, Okamoto Y, Yoshioka K, Takuwa N (2012) Sphingosine-1-phosphate signaling in physiology and diseases. Biofactors 38(5):329–337.

48. Arish M, Alaidarous M, Ali R, Akhter Y, Rub A (2017) Implication of sphingosine-1-phosphate signaling in diseases: molecular mechanism and therapeutic strategies. J Recept Signal Transduct Res 37(5):437–446.

49. Rose DM, Alon R, Ginsberg MH (2007) Integrin modulation and signaling in leukocyte adhesion and migration. Immunol Rev 218:126–134.

50. Abram CL, Lowell CA (2009) The ins and outs of leukocyte integrin signaling. Annu Rev Immunol 27:339–362.

51. Luo B-H, Carman CV, Springer TA (2007) Structural basis of integrin regulation and signaling. Annu Rev Immunol 25:619–647.

52. Bendall SC, Nolan GP, Roederer M, Chattopadhyay PK (2012) A deep profiler’s guide to cytometry. Trends Immunol 33(7):323–332.

53. Terfve CDA, Wilkes EH, Casado P, Cutillas PR, Saez-Rodriguez J (2015) Large-scale models of signal propagation in human cells derived from discovery phosphoproteomic data. Nat Commun 6:8033.

54. Bulusu KC, et al. (2016) Modelling of compound combination effects and applications to efficacy and toxicity: state-of-the-art, challenges and perspectives. Drug Discov Today 21(2):225–238.

55. Halasz M, Kholodenko BN, Kolch W, Santra T (2016) Integrating network reconstruction with mechanistic modeling to predict cancer therapies. Sci Signal 9(455):ra114.

56. Martinez-Pasamar S, et al. (2013) Dynamic cross-regulation of antigen-specific effector and regulatory T cell subpopulations and microglia in brain autoimmunity. BMC Syst Biol 7:34.

