## Supplementary Figures S1-S5 for "Prediction of combination therapies based on topological modeling of the immune signaling network in Multiple Sclerosis"

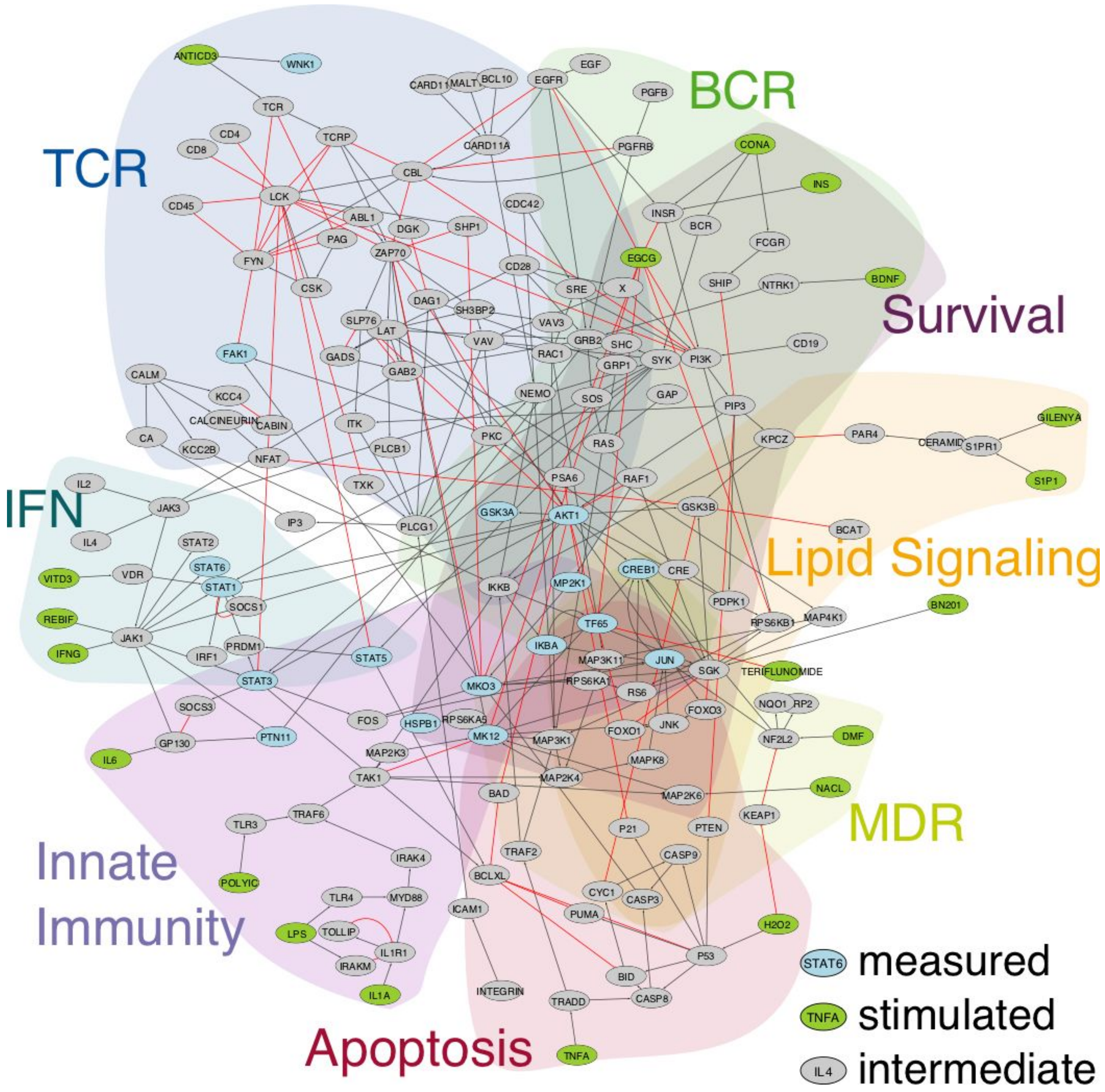


**Supplementary Figure S1. Reference-based curated MS- and immune-specific Prior Knowledge signaling Network (PKN).** The shadows indicate the signaling pathways studied (interferon response (IFNβ), B-cell receptor (BCR) signaling, T- cell receptor (TCR) signaling, cellular survival and apoptosis, lipid signaling, innate immunity and multi-drug response (MDR) genes), including the crosstalk among them. Blue ovals: experimentally measured phosphoproteins; grey ovals: non-measured phosphoproteins and other molecules involved in signaling; green ovals: stimuli used in the *in vitro* assays; red lines: inhibitory reactions; black lines: activatory reactions.

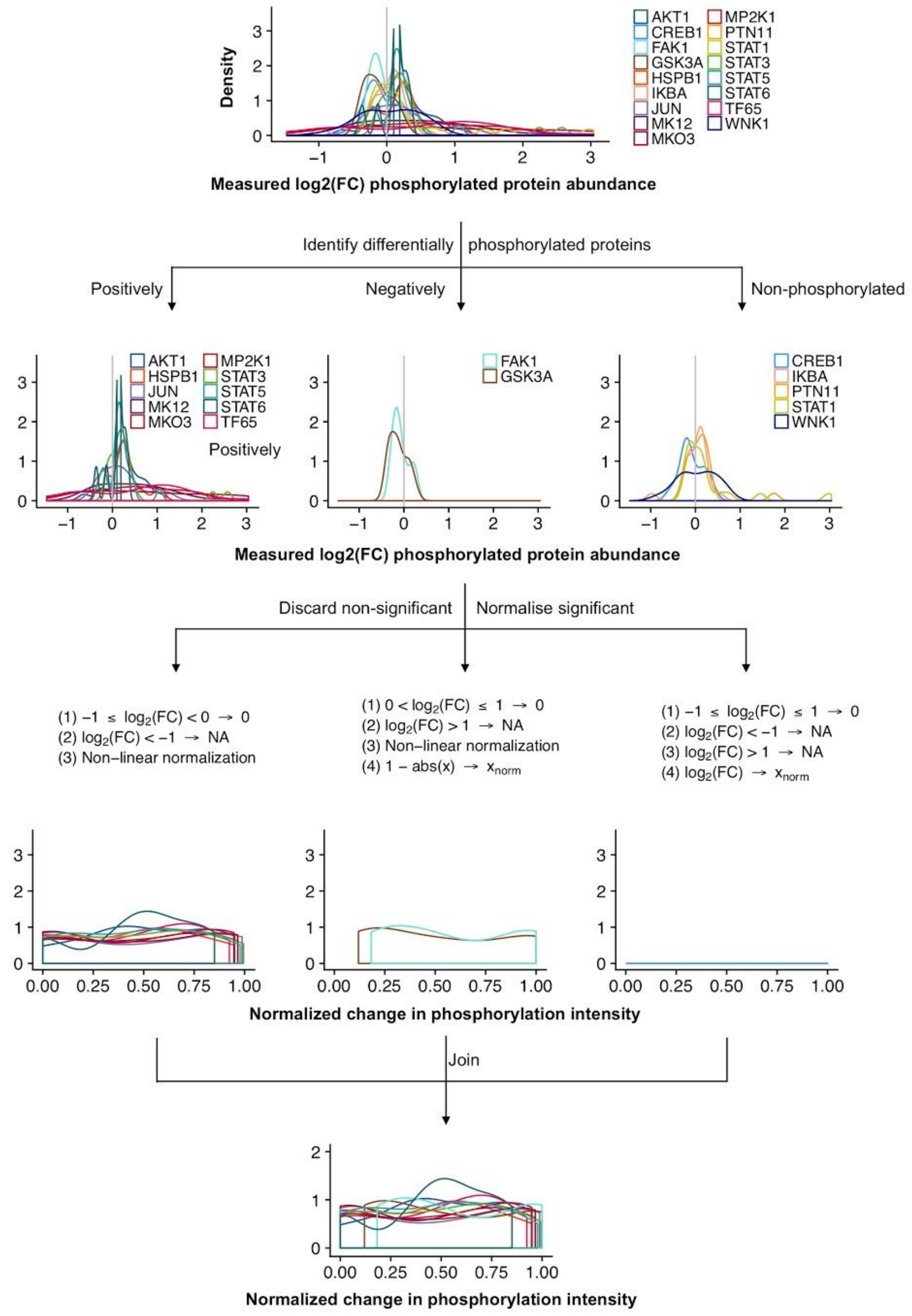
Density
AKT1
CREB1
FAK1
GSK3A
HSPB1
IKBA
JUN
MK12
MKO3
MP2K1
PTN11
STAT1
STAT3
STAT5
STAT6
TF65
WNK1
Measured log2(FC) phosphorylated protein abundance
Identify differentially phosphorylated proteins
Positively
Negatively
Non-phosphorylated
Positively
Measured log2(FC) phosphorylated protein abundance
Discard non-significant
Normalise significant
(1) −1 ≤ log2(FC) < 0 → 0
(2) log2(FC) < −1 → NA
(3) Non−linear normalization
(1) 0 < log2(FC) ≤ 1 → 0
(2) log2(FC) > 1 → NA
(3) Non−linear normalization
(4) 1 − abs(x) → xnorm
(1) −1 ≤ log2(FC) ≤ 1 → 0
(2) log2(FC) < −1 → NA
(3) log2(FC) > 1 → NA
(4) log2(FC) → xnorm
Normalized change in phosphorylation intensity
Join
Normalized change in phosphorylation intensity

**Supplementary Figure S2. Phosphoproteomics normalization pipeline**. The graph shows the normalization algorithm developed to identify and select the significantly positive and negative phosphorylated measurements, and normalize them accordingly to allow Boolean modeling (see Methods). For consistency, the data highlighted in main Figure 2 (patient KI044) was used to visualize the normalization process.

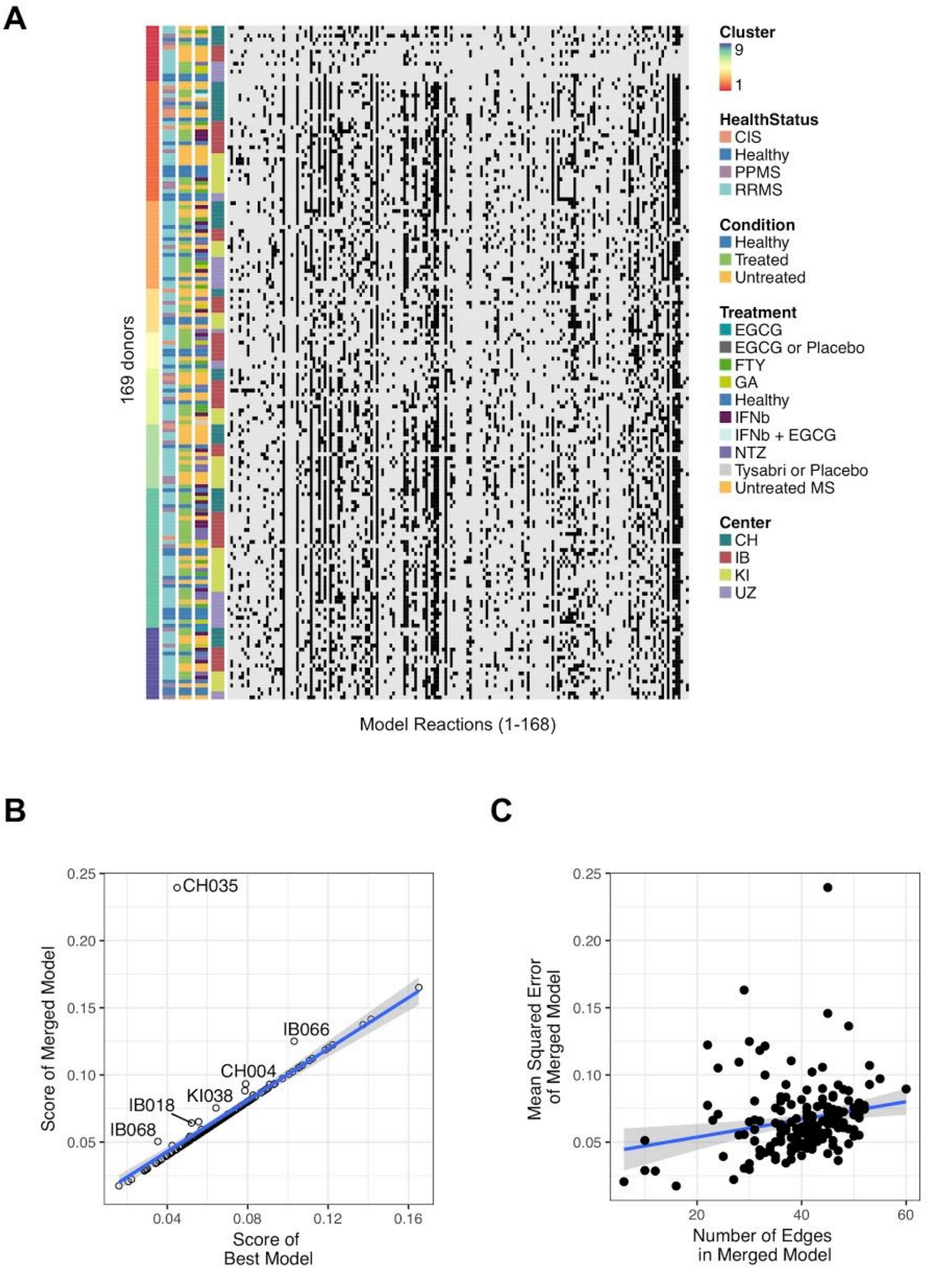
A
169 donors
Model Reactions (1-168)
Cluster
9
1
HealthStatus
CIS
Healthy
PPMS
RRMS
Condition
Healthy
Treated
Untreated
Treatment
EGCG
EGCG or Placebo
FTY
GA
Healthy
IFNb
IFNb + EGCG
NTZ
Tysabri or Placebo
Untreated MS
Center
CH
IB
KI
UZ
B
Score of Merged Model
Score of Best Model
CH035
IB066
CH004
KI038
IB018
IB068
0.25
0.20
0.15
0.10
0.05
0.04
0.08
0.12
0.16
C
Mean Squared Error of Merged Model
Number of Edges in Merged Model
20
40
60

**Supplementary Figure S3. Model quality analysis confirms robustness at the single patient level. A)** Signaling network found by modeling for each donor, visualized as a heatmap. Rows: Single donor network. Columns: Signaling activity determined for each interaction by calibrating the PKN shown in Supplementary Figure S1 after removing the unidentifiable interactions using the phosphoproteomics dataset of each donor. No bias was found due to confounding variables using affinity propagation clustering of the final donor models. The resulting clusters are not enriched for treatment, center, disease subtype or medical condition; **B)** Score of best model found for each patient compared to the final median solution. No new, median models were found to be better than their corresponding best solution, indicating successful optimization; **C)** No relationship was found between model size and performance as quantified by Mean Squared Error between model simulation and data, supporting model quality.

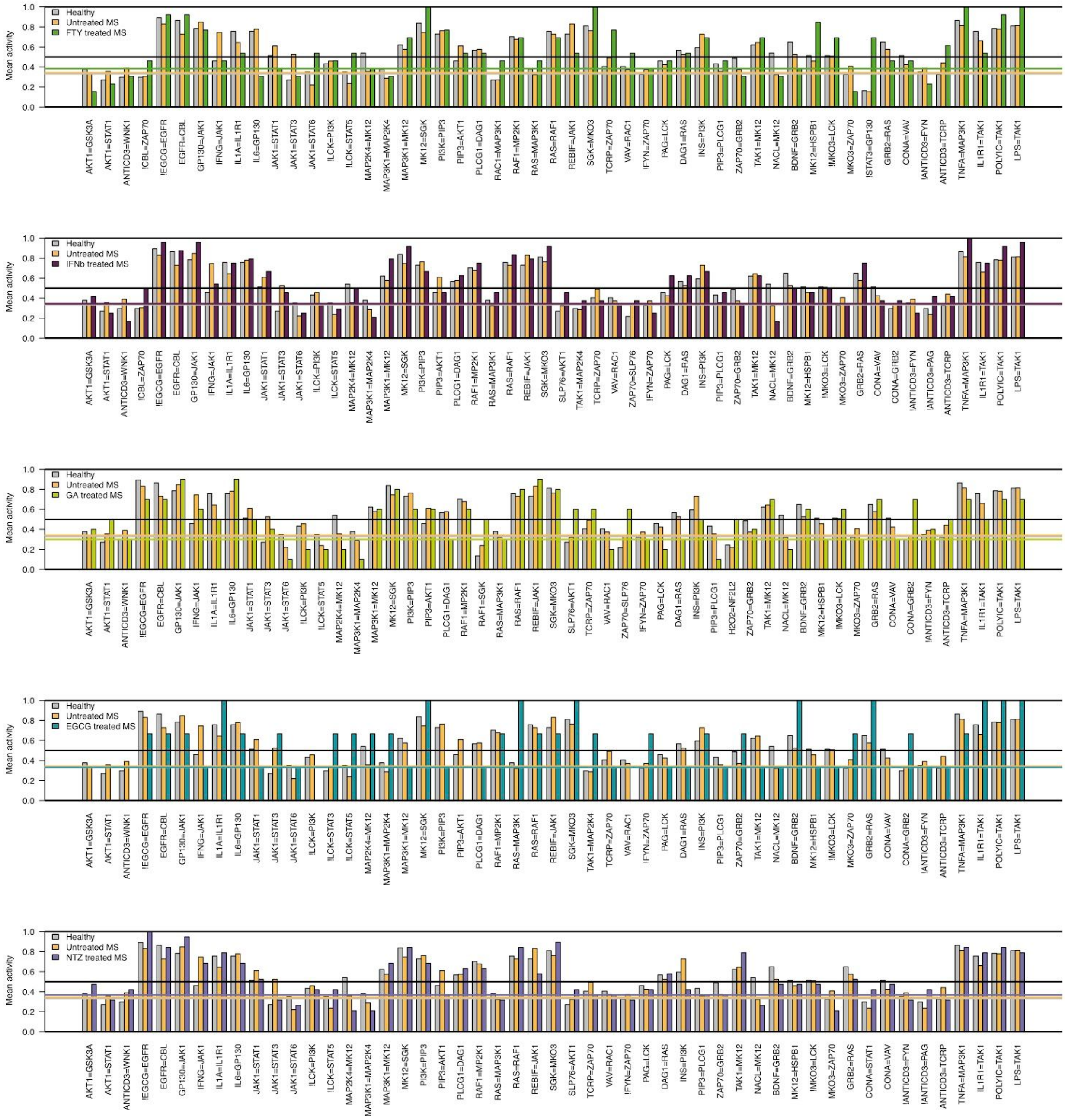


**Supplementary Figure S4. Signaling activity of each interaction of the subgroup networks.** Bars represent the signaling activity of each interaction for each donor subgroup, after merging the models for each individual donor by the mean across subgroup donors. Each one of the 5 panels shows the signaling activity of each interaction of one of the 5 drug treatment subgroups against the activity in healthy donors (grey) and untreated patients (orange). The colored thresholds show the upper quartile of each corresponding subgroup mean defining as signaling noise those interactions with signaling activity below that threshold.

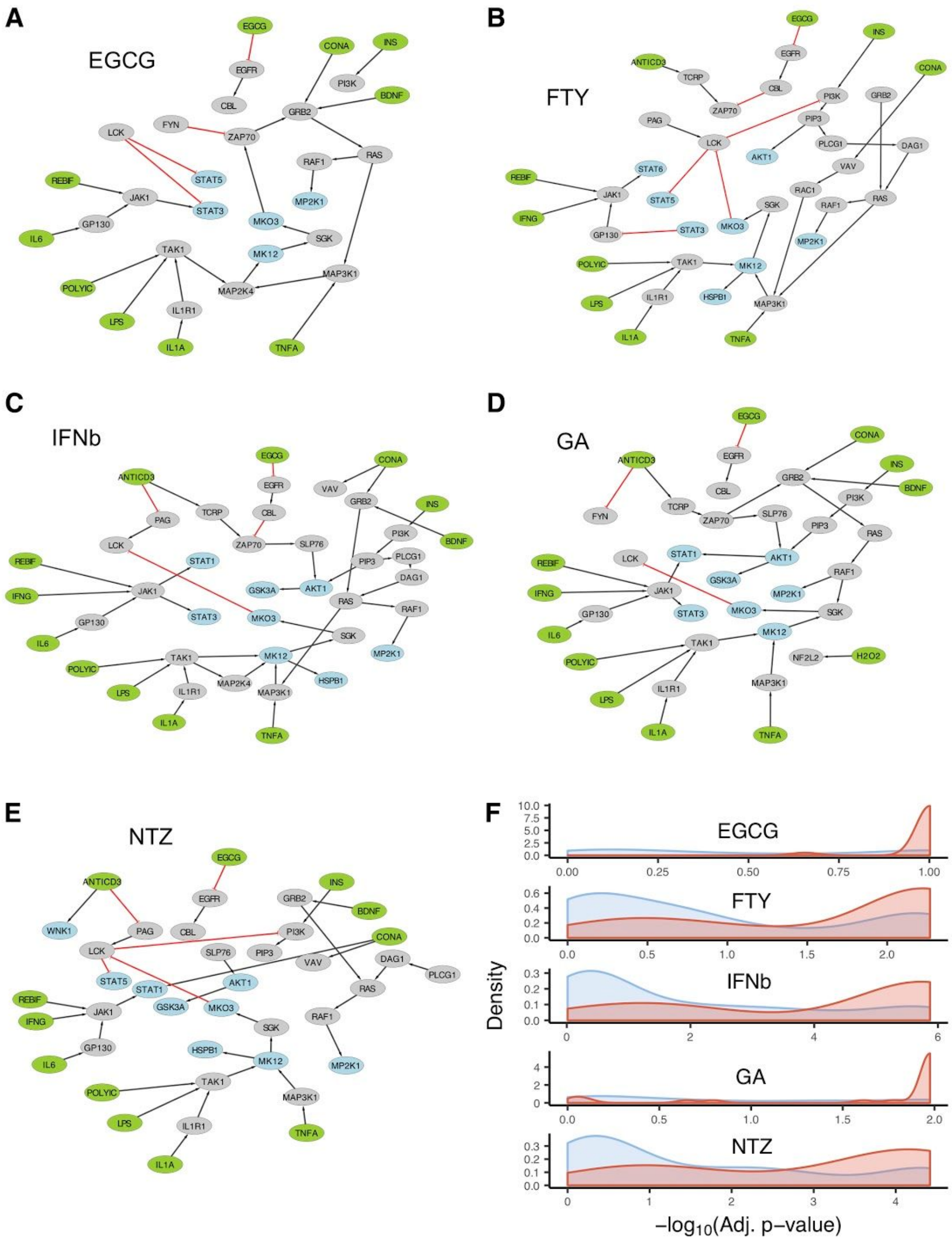
A
EGCG
B
FTY
C
IFNb
D
GA
E
NTZ
F
EGCG
FTY
IFNb
GA
NTZ
Density
−log10(Adj. p−value)

**Supplementary Figure S5. Signaling networks found for each MS-specific therapy. A-E)** The models previously calculated for each patient were merged to reveal the common active pathways for the experimental drug EGCG and for each MS drug (FTY, IFNβ, GA and NTZ); **F)** The differentially phosphorylated proteins were overrepresented (x axis shows the –log10 adjusted p-value) in pathways predicted by modeling as a density score (red line: protein found in model for each corresponding therapy, blue line: absent in that model) and confirmed statistically (see main text).
