## Supplementary Tables S1-S8 for "Prediction of combination therapies based on topological modeling of the immune signaling network in Multiple Sclerosis"

**Supplementary Table S1** References of the kinase interactions of the Prior Knowledge Network (PKN)

| No | Interaction | Reference |
|---|---|---|
| 1 | ABL1⟶ZAP70 | Saez-Rodriguez2007, Zipfel2004 |
| 2 | AKT1⟞BAD | Saez-Rodriguez2007, Hanada2004, |
| 3 | AKT1⟶CREB1 | Du1998 |
| 4 | AKT1⟞FOXO1 | Saez-Rodriguez2007, Hanada2004 |
| 5 | AKT1⟶FOXO3 | Brunet2001 |
| 6 | AKT1⟶GSK3A | Saez-Rodriguez2007, Hanada2004, Liang2003 |
| 7 | AKT1⟶GSK3B | Saez-Rodriguez2007, Hanada2004, Liang2003 |
| 8 | AKT1⟶IKKB | Dan2008 |
| 9 | AKT1⟞P21 | Saez-Rodriguez2007, Hanada2004, Liang2003 |
| 10 | AKT1⟞PSA6 | Saez-Rodriguez2007, Hanada2004, Liang2003 |
| 11 | AKT1⟞RAF1 | Moelling2002 |
| 12 | AKT1⟞SLP76 | |
| 13 | AKT1⟶STAT1 | Beyer2011 |
| 14 | AKT1⟶STAT3 | Beyer2011 |
| 15 | AKT1⟞ZAP70 | |
| 16 | ANTICD3⟶TCR | Model Input |
| 17 | ANTICD3⟶WNK1 | Mayya2009 |
| 18 | BAD⟞BCLXL | Saez-Rodriguez2007, Zha1997, Yang1995 |
| 19 | BCL10⟶CARD11A | Saez-Rodriguez2007, Thome2004, Weil2006, Hayden2004 |
| 20 | BCLXL⟞BID | Hengartner2000 |
| 21 | BCLXL⟞P53 | Levine2006 |
| 22 | BCR⟶SYK | Gold2002 |
| 23 | BDNF⟶NTRK1 | |
| 24 | BID⟶CYC1 | Hengartner2000 |
| 25 | BN201⟶SGK | |
| 26 | CA⟶CALM | Saez-Rodriguez2007, Feske2001 |
| 27 | CABIN⟞CALCINEURIN | Saez-Rodriguez2007, Matsuda2000, Ryeom2003, Kashashian1998 |
| 28 | CALCINEURIN⟶NFAT | Saez-Rodriguez2007, Huang2004, Macian2005, Krauss2006 |
| 29 | CALM⟶CALCINEURIN | Saez-Rodriguez2007, Matsuda2000, Ryeom2003, |

| | | |
|---|---|---|
| | | Kashashian1998 |
| 30 | CALM⟶KCC2B | Saez-Rodriguez2007, Hughes2001 |
| 31 | CALM⟶KCC4 | Saez-Rodriguez2007, Anderson1998 |
| 32 | CARD11⟶CARD11A | Saez-Rodriguez2007, Thome2004, Weil2006, Hayden2004 |
| 33 | CARD11A⟶NEMO | Saez-Rodriguez2007, Thome2004, Weil2006, Hayden2004 |
| 34 | CASP3⟶FAK1 | Schlaepfer1997 |
| 35 | CASP8⟶BID | Hengartner2000 |
| 36 | CASP8⟶CASP3 | Hengartner2000 |
| 37 | CASP9⟶CASP3 | Hengartner2000 |
| 38 | CBL⟞EGFR | Galisteo1995, Tarcic2009 |
| 39 | CBL⟶FYN | Duan2004, Kaabeche2004 |
| 40 | CBL⟶LCK | Duan2004 |
| 41 | CBL⟞PGFRB | Reddi2007, Wardega2010 |
| 42 | CBL⟞PI3K | Saez-Rodriguez2007, Fang2001 |
| 43 | CBL⟞TCRP | Saez-Rodriguez2007, Huang2004, Rao2000 |
| 44 | CBL⟞ZAP70 | Saez-Rodriguez2007, Zipfel2004, Rao2000 |
| 45 | CD19⟶PI3K | Gold2002 |
| 46 | CD28⟶GADS | Ellis2000 |
| 47 | CD28⟶GRB2 | Okkenhaug1998, Nunes1996 |
| 48 | CD28⟶PI3K | Saez-Rodriguez2007, August1994, Ghiotto-Ragueneau1996 |
| 49 | CD28⟶X | Saez-Rodriguez2007 |
| 50 | CD4⟞LCK | Saez-Rodriguez2007, Palacios2004 |
| 51 | CD45⟞FYN | Saez-Rodriguez2007, Filipp2004a, Palacios2004 |
| 52 | CD45⟞LCK | Saez-Rodriguez2007, Palacios2004 |
| 53 | CD8⟞LCK | Palacios2004 |
| 54 | CDC42⟶MAP3K1 | Saez-Rodriguez2007, Fanger1997, Kaga1998 |
| 55 | CDC42⟶SRE | Saez-Rodriguez2007, Hill1995 |
| 56 | CERAMIDE⟶PAR4 | Wang2005 |
| 57 | CONA⟶BCR | DePetris1975 |

| | | |
|---|---|---|
| 58 | CONA⟶FCGR | DePetris1975 |
| 59 | CONA⟶INSR | Cuatrecasas1973 |
| 60 | CREB1⟶CRE | Saez-Rodriguez2007, Krauss2003 |
| 61 | CREB1⟶JUN | |
| 62 | CSK⟶FYN | Palacios2004 |
| 63 | CSK⟶LCK | Saez-Rodriguez2007, Palacios2004 |
| 64 | CYC1⟶CASP9 | Hengartner2000 |
| 65 | DAG1⟶GRP1 | Saez-Rodriguez2007, DiFiore2003, Bivona2003 |
| 66 | DAG1⟶PKC | Saez-Rodriguez2007, Lee2005 |
| 67 | DGK⟞DAG1 | Saez-Rodriguez2007, Topham2006 |
| 68 | DMF⟶NF2L2 | Gold2012 |
| 69 | EGCG⟞EGFR | Michailidou2015, Gupta2010, VanAller2011 |
| 70 | EGCG⟞INSR | |
| 71 | EGCG⟞MK12 | Michailidou2015, Gupta2010, VanAller2011 |
| 72 | EGCG⟞MK03 | Michailidou2015, Gupta2010, VanAller2011 |
| 73 | EGCG⟞PI3K | Michailidou2015, Gupta2010, VanAller2011 |
| 74 | EGCG⟞RPS6KB1 | |
| 75 | EGCG⟞TF65 | Michailidou2015, Gupta2010 |
| 76 | EGF⟶EGFR | Galisteo1995 |
| 77 | EGFR⟶CBL | Galisteo1995 |
| 78 | EGFR⟶GRB2 | Galisteo1995 |
| 79 | EGFR⟶INSR | |
| 80 | FCGR⟶SHIP | Gold2002 |
| 81 | FOS⟶JUN | Saez-Rodgriguez2007, Huang2004, Krauss2003 |
| 82 | FYN⟞ABL1 | Saez-Rodriguez2007, Zipfel2004 |
| 83 | FYN⟞FAK1 | Messina2003, Arold2001 |
| 84 | FYN⟞PAG | Saez-Rodriguez2007, Davidson2003 |
| 85 | FYN⟞TCRP | Saez-Rodriguez2007, Filipp2004 |
| 86 | GAB2⟞SLP76 | Saez-Rodriguez2007, Yamasaki2001, Yamasaki2003 |
| 87 | GADS⟶GAB2 | Saez-Rodriguez2007, Yamasaki2001, Yamasaki2003 |
| 88 | GADS⟶SLP76 | Saez-Rodriguez2007, |

| | | |
|---|---|---|
| | | Horejsi2004, Togni2004 |
| 89 | GAP⟶RAS | Saez-Rodriguez2007, Genot2000 |
| 90 | GILENYA⟶S1PR1 | |
| 91 | GP130⟶JAK1 | Heinrich1999, Murray2007 |
| 92 | GP130⟶PTN11 | Schaper1998 |
| 93 | GRB2⟶GAB2 | Saez-Rodriguez2007, Yamasaki2001, Yamasaki2003 |
| 94 | GRB2⟶SOS | Saez-Rodriguez2007, Buday1994 |
| 95 | GRP1⟶RAS | Saez-Rodriguez2007, DiFiore2003 |
| 96 | GSK3B⟞BCAT | Saez-Rodriguez2007, Liang2003 |
| 97 | GSK3B⟞CYC1 | Saez-Rodriguez2007, Liang2003 |
| 98 | GSK3B⟞NFAT | |
| 99 | H2O2⟞KEAP1 | Schulze2012 |
| 100 | H2O2⟶P53 | Hofseth2004 |
| 101 | ICAM1⟶PLCG1 | Etienne-Manneville2000 |
| 102 | IFNG⟶JAK1 | Pertsovskaya2013 |
| 103 | IKBA⟶TF65 | Saez-Rodriguez2007, Huang2004, Mattioli2004 |
| 104 | IKKB⟶IKBA | Saez-Rodriguez2007, Huang2004, Krauss2003 |
| 105 | IL1A⟶IL1R1 | Weber2010 |
| 106 | IL1R1⟶MYD88 | Weber2010 |
| 107 | IL1R1⟶TOLLIP | Weber2010 |
| 108 | IL2⟶JAK3 | Kelly-Welch2005 |
| 109 | IL4⟶JAK3 | Kelly-Welch2005 |
| 110 | IL6⟶GP130 | Hirano1998 |
| 111 | INS⟶INSR | Saltiel2001 |
| 112 | INSR⟶PI3K | Saltiel2001 |
| 113 | INTEGRIN⟶ICAM1 | Etienne-Manneville2000 |
| 114 | IP3⟶CA | Saez-Rodriguez2007, Chan1999 |
| 115 | IRAK4⟶TRAF6 | Weber2010 |
| 116 | IRAKM⟞IL1R1 | Weber2010 |
| 117 | ITK⟶PLCG1 | Saez-Rodriguez2007, Czar2001, Rellahan2003 |
| 118 | JAK1⟶AKT1 | Beyer2011 |
| 119 | JAK1⟶PTN11 | Schaper1998 |
| 120 | JAK1⟶STAT1 | Murray2007 |
| 121 | JAK1⟶STAT2 | Kisseleva2002 |

| | | |
|---|---|---|
| 122 | JAK1⟶STAT3 | Beyer2011, Heinrich1999, Murray2007 |
| 123 | JAK1⟶STAT6 | Kelly-Welch2005, Murray2007 |
| 124 | JAK3⟶GAB2 | Beyer2011 |
| 125 | JAK3⟶JAK1 | Beyer2011 |
| 126 | JAK3⟶SYK | Kelly-Welch2005, Beyer2011 |
| 127 | JNK⟶FOXO1 | Huang2007 |
| 128 | JNK⟶FOXO3 | Huang2007 |
| 129 | JNK⟶JUN | Saez-Rodriguez2007, Krauss2003 |
| 130 | JUN⟶CREB1 | |
| 131 | KCC2B⟶IKKB | Saez-Rodriguez2007, Thome2004, Weil2006, Hayden2004 |
| 132 | KCC4⊣CABIN | Saez-Rodriguez2007, Pan2005 |
| 133 | KEAP1⊣NF2L2 | Schulze2012 |
| 134 | KPCZ⟶GSK3B | Wang2005 |
| 135 | KPCZ⟶IKKB | Wang2005 |
| 136 | LAT⟶GAB2 | Saez-Rodriguez2007, Yamasaki2001, Yamasaki2003 |
| 137 | LAT⟶GADS | Saez-Rodriguez2007, Horejsi2004, Togni2004 |
| 138 | LAT⟶GRB2 | Saez-Rodriguez2007, Lindquist2003, Horejsi2004 |
| 139 | LAT⟶MAP4K1 | Saez-Rodriguez2007, Liou2000 |
| 140 | LAT⟶PLCB1 | Saez-Rodriguez2007, Horejsi2004, Togni2004 |
| 141 | LAT⟶SH3BP2 | Saez-Rodriguez2007, Qu2005 |
| 142 | LCK⊣ABL1 | Saez-Rodriguez2007, Zipfel2004 |
| 143 | LCK⟶FYN | Saez-Rodriguez2007, Zamoyska2003, Filipp2004a |
| 144 | LCK⊣PI3K | Saez-Rodriguez2007, Deane2004 |
| 145 | LCK⊣STAT3 | Beyer2011 |
| 146 | LCK⊣STAT5 | Beyer2011 |
| 147 | LCK⊣TCRP | Saez-Rodriguez2007, Zamoyska2003, Filipp2004a |
| 148 | LCK⊣TXK | Saez-Rodriguez2007, Shan2000 |
| 149 | LCK⊣ZAP70 | Palacios2004 |
| 150 | LPS⟶IRAKM | Weber2010 |
| 151 | LPS⟶TLR4 | Chastain2012 |
| 152 | MALT1⟶CARD11A | Saez-Rodriguez2007, Thome2004, Weil2006, Hayden2004 |

| | | |
|---|---|---|
| 153 | MAP2K3⟶MK12 | |
| 154 | MAP2K4⟶MAPK8 | |
| 155 | MAP2K4⟶MK12 | |
| 156 | MAP2K6⟶MK12 | Raingeaud1996 |
| 157 | MAP3K1⟶MAP2K4 | Saez-Rodriguez2007, Yan1994 |
| 158 | MAP3K1⟶MK12 | Saez-Rodriguez2007, Guan1998 |
| 159 | MAP3K11⟶MAP2K4 | Saez-Rodriguez2007, Tibbles1996 |
| 160 | MAP4K1⟶MAP3K1 | Saez-Rodriguez2007, Hu1996 |
| 161 | MAP4K1⟶MAP3K11 | Saez-Rodriguez2007, Tibbles1996 |
| 162 | MAPK8⟶JUN | |
| 163 | MK12⟶CREB1 | |
| 164 | MK12⟶JNK | Saez-Rodriguez2007, Davis2000 |
| 165 | MK12⟶RPS6KA5 | |
| 166 | MK12⟶SGK | BelAiba2006 |
| 167 | MK12—⊣TAK1 | Weber2010 |
| 168 | MK03⟶FOS | Saez-Rodriguez2007, Huang2004 |
| 169 | MK03⟶JUN | |
| 170 | MK03⟶RPS6KA1 | Saez-Rodriguez2007, Frdin1999 |
| 171 | MK03—⊣SHP1 | Saez-Rodriguez2007 |
| 172 | MK03—⊣ZAP70 | |
| 173 | MP2K1⟶MK03 | Saez-Rodriguez2007, Huang2004, Krauss2003 |
| 174 | MYD88⟶IRAK4 | Weber2010, Kawai2010 |
| 175 | NACL⟶MAP2K6 | Kleinewietfeld2013 |
| 176 | NEMO⟶IKKB | Saez-Rodriguez2007, Thome2004, Weil2006, Hayden2004 |
| 177 | NF2L2⟶CREB1 | Katoh2001 |
| 178 | NF2L2⟶JUN | Venugopal1998 |
| 179 | NF2L2⟶MRP2 | Maher2007 |
| 180 | NF2L2⟶NQO1 | Maher2007 |
| 181 | NTRK1⟶GRB2 | |
| 182 | P53⟶BID | Levine2006 |
| 183 | P53⟶CASP8 | Levine2006 |
| 184 | P53⟶CASP9 | Levine2006 |
| 185 | P53⟶P21 | Watcharasit2002 |
| 186 | P53⟶PTEN | Levine2006 |

| | | |
|---|---|---|
| 187 | P53⟶PUMA | Madapura2012 |
| 188 | PAG⟶CSK | Saez-Rodriguez2007, Lindquist2003, Horejsi2004 |
| 189 | PAR4⟞KPCZ | Wang2005 |
| 190 | PDPK1⟶AKT1 | Saez-Rodriguez2007, Lafont2000, Alessi1997, Anderson1998 |
| 191 | PDPK1⟶PKC | Saez-Rodriguez2007, Lee2005 |
| 192 | PDPK1⟶RPS6KB1 | Saez-Rodriguez2007, Pullen1998, Hinton2004 |
| 193 | PGFB⟶PGFRB | |
| 194 | PGFRB⟶CBL | |
| 195 | PGFRB⟶GRB2 | |
| 196 | PI3K⟶PIP3 | Saez-Rodriguez2007, Okkenhaug2004, Rameh1999 |
| 197 | PI3K⟶RPS6KB1 | |
| 198 | PIP3⟶AKT1 | Song2005 |
| 199 | PIP3⟶ITK | Saez-Rodriguez2007, Huang2004, Togni2004, Czar2001 |
| 200 | PIP3⟶KPCZ | Beyer2011 |
| 201 | PIP3⟶PDPK1 | Saez-Rodriguez2007, Mora2004 |
| 202 | PKC⟶FAK1 | Etienne-Manneville2000 |
| 203 | PKC⟶NEMO | Saez-Rodriguez2007, Koshnan2000 |
| 204 | PLCB1⟶PLCG1 | Saez-Rodriguez2007, Czar2001, Rellahan2003 |
| 205 | PLCG1⟶DAG1 | Saez-Rodriguez2007, Huang2004 |
| 206 | PLCG1⟶IP3 | Saez-Rodriguez2007, Huang2004, Togni2004 |
| 207 | PLCG1⟶PKC | Etienne-Manneville2000 |
| 208 | POLYIC⟶TLR3 | Bsibsi2012 |
| 209 | PTEN⟞PIP3 | Saez-Rodriguez2007, Okkenhaug2004, Rameh1999 |
| 210 | PTN11⟶GRB2 | Schaper1998, Kim1999 |
| 211 | PUMA⟞BCLXL | Madapura2012 |
| 212 | RAC1⟶MAP3K1 | Saez-Rodriguez2007, Fanger1997 |
| 213 | RAC1⟶MAP3K11 | Saez-Rodriguez2007, Teramoto1996 |
| 214 | RAC1⟶SRE | Saez-Rodriguez2007, Hill1995 |
| 215 | RAF1⟶MP2K1 | Saez-Rodriguez2007, Franklin1994 |
| 216 | RAF1⟶SGK | |
| 217 | RAS⟶MAP3K1 | |

| | | |
|---|---|---|
| 218 | RAS⟶RAF1 | Saez-Rodriguez2007, Krauss2003 |
| 219 | REBIF⟶JAK1 | Pertsovskaya2013 |
| 220 | RPS6KA1⟶RS6 | Saez-Rodriguez2007, Frdin1999 |
| 221 | RPS6KA5⟶HSPB1 | |
| 222 | RPS6KB1⟶RS6 | Saez-Rodriguez2007, Frdin1999 |
| 223 | RS6⟶CREB1 | Saez-Rodriguez2007, Frdin1999 |
| 224 | S1P1⟶S1PR1 | Lee1998 |
| 225 | S1PR1⟶CERAMIDE | |
| 226 | SGK⟶CREB1 | David2005 |
| 227 | SGK⟞FOXO1 | Feroze-Zaidi2007 |
| 228 | SGK⟞FOXO3 | Brunet2001 |
| 229 | SGK⟶GSK3B | Aoyama2005 |
| 230 | SGK⟶IKBA | Zhang2005 |
| 231 | SGK⟶IKKB | Zhang2005 |
| 232 | SGK⟶MK03 | |
| 233 | SGK⟶RPS6KB1 | Biondi2001 |
| 234 | SH3BP2⟶VAV | Saez-Rodriguez2007, Qu2005 |
| 235 | SH3BP2⟶VAV3 | Saez-Rodriguez2007, Zakaria2004 |
| 236 | SHC⟶GRB2 | Beyer2011 |
| 237 | SHIP⟞PIP3 | Saez-Rodriguez2007, Okkenhaug2004, Rameh1999 |
| 238 | SHP1⟶LCK | Saez-Rodriguez2007, Palacios2004 |
| 239 | SHP1⟞ZAP70 | Alegre2001 |
| 240 | SLP76⟶AKT1 | |
| 241 | SLP76⟶ITK | Saez-Rodriguez2007, Huang2004, Togni2004, Czar2001 |
| 242 | SLP76⟶VAV | Huang2004 |
| 243 | SOCS1⟞STAT1 | Chen2000 |
| 244 | SOCS3⟞GP130 | Zhang2006 |
| 245 | SOS⟶RAS | Saez-Rodriguez2007, DiFiore2003 |
| 246 | STAT1⟶IRF1 | Pertsovskaya2013 |
| 247 | STAT1⟶PRDM1 | Beyer2011 |
| 248 | STAT1⟶SOCS1 | Chen2000 |
| 249 | STAT3⟶BCLXL | Kisseleva2002 |
| 250 | STAT3⟶FOS | Yang2003 |
| 251 | STAT3⟶PRDM1 | Beyer2011 |

| | | |
|---|---|---|
| 252 | STAT3⟶SOCS3 | Zhang2006 |
| 253 | STAT5⟶BCLXL | Murray2007 |
| 254 | STAT5⟶PRDM1 | Beyer2011 |
| 255 | SYK⟶PLCG1 | Gold2002 |
| 256 | SYK⟶SHC | Maghazachi2005 |
| 257 | SYK⟶STAT1 | Beyer2011 |
| 258 | SYK⟶STAT3 | Beyer2011 |
| 259 | SYK⟶STAT5 | Beyer2011 |
| 260 | SYK⟶VAV | Gold2002 |
| 261 | TAK1⟶IKKB | Weber2010 |
| 262 | TAK1⟶MAP2K3 | Weber2010 |
| 263 | TAK1⟶MAP2K4 | Weber2010 |
| 264 | TAK1⟶MAP2K6 | Weber2010 |
| 265 | TCR⟞FYN | Saez-Rodriguez2007, Filipp2004 |
| 266 | TCR⟞PAG | Saez-Rodriguez2007, Davidson2003 |
| 267 | TCR⟶TCRP | Saez-Rodriguez2007, Filipp2004a |
| 268 | TCRP⟶DGK | Saez-Rodriguez2007, Sanjuan2001 |
| 269 | TCRP⟶ZAP70 | Saez-Rodriguez2007, Zipfel2004 |
| 270 | TERIFLUNOMIDE⟞TF65 | Manna1999 |
| 271 | TF65⟶IKBA | Weber2010 |
| 272 | TLR3⟶TRAF6 | Kawai2010 |
| 273 | TLR4⟶MYD88 | Kawai2010 |
| 274 | TNFA⟶TRADD | Naude2011, Newton2012 |
| 275 | TOLLIP⟞IL1R1 | Weber2010 |
| 276 | TRADD⟶CASP8 | Naude2011, Newton2012 |
| 277 | TRADD⟶TRAF2 | Martinon2005 |
| 278 | TRAF2⟶IKKB | Martinon2005, Newton2012 |
| 279 | TRAF2⟶MAP3K1 | Newton2012 |
| 280 | TRAF6⟶TAK1 | Weber2010 |
| 281 | TXK⟶PLCG1 | Saez-Rodriguez2007, Czar2001, Rellahan2003 |
| 282 | VAV⟶PKC | Saez-Rodriguez2007, Villabala2002 |
| 283 | VAV⟶PLCG1 | Saez-Rodriguez2007, Czar2001, Rellahan2003 |
| 284 | VAV⟶RAC1 | Saez-Rodriguez2007, Zakaria2004 |
| 285 | VAV3⟶RAC1 | Saez-Rodriguez2007, |

| | | |
|---|---|---|
| | | Zakaria2004 |
| 286 | VDR⟶STAT1 | Vidal 2002 |
| 287 | VITD3⟶VDR | Vidal 2002 |
| 288 | X⟶PI3K | Saez-Rodriguez2007 |
| 289 | X⟶VAV | Saez-Rodriguez2007 |
| 290 | ZAP70⟶GAB2 | Saez-Rodriguez2007, Togni2004, Yamasaki2001, Yamasaki2003 |
| 291 | ZAP70⟶LAT | Saez-Rodriguez2007, Huang2004 |
| 292 | ZAP70⟶MK12 | Saez-Rodriguez2007, Salvador2005 |
| 293 | ZAP70⟶SH3BP2 | Saez-Rodriguez2007, Qu2005 |
| 294 | ZAP70⟶SLP76 | Saez-Rodriguez2007, Horejsi2004, Togni2004 |

### Supplementary Table S2

List of the phosphoproteins selected from the literature search, the stimuli applied and the MS drugs used for building the Prior Knowledge Network (PKN)

| No | Model Name | Name | UniProt-ID / ChEMBL-ID |
|---|---|---|---|
| 1 | ABL1 | Tyrosine-protein kinase ABL1 | P00519 |
| 2 | AKT1 | RAC-alpha serine/threonine-protein kinase | P31749 |
| 3 | ANTICD3 | Antibody against human CD3 | |
| 4 | BAD | Bcl2-associated agonist of cell death | Q92934 |
| 5 | BCAT | Catenin beta-1 | P35222 |
| 6 | BCL10 | B-cell lymphoma/leukemia 10 | O95999 |
| 7 | BCLXL | Bcl-2-like protein 1 | Q07817 |
| 8 | BCR | B-cell receptor | |
| 9 | BDNF | Brain-derived neurotrophic factor | P23560 |
| 10 | BID | BH3-interacting domain death agonist | P55957 |
| 11 | BN201 | Drug candidate by Bionure | |
| 12 | CA | Calcium | CHEMBL2146121 |
| 13 | CABIN | Calcineurin-binding protein cabin-1 | Q9Y6J0 |
| 14 | CALCINEURIN | Calcineurin phosphatase family | |
| 15 | CALM | Calmodulin | P62158 |
| 16 | CARD11 | Caspase recruitment domain-containing protein 11 | Q9BXL7 |
| 17 | CARD11A | Complex of BCL10, CARD11, and MALT1 | |
| 18 | CASP3 | Caspase-3 | P42574 |
| 19 | CASP8 | Caspase-8 | Q14790 |
| 20 | CASP9 | Caspase-9 | P55211 |
| 21 | CBL | E3 ubiquitin-protein ligase CBL | P22681 |
| 22 | CD19 | B-lymphocyte antigen CD19 | P15391 |
| 23 | CD28 | T-cell-specific surface glycoprotein CD28 | P10747 |
| 24 | CD4 | T-cell surface glycoprotein CD4 | P01730 |
| 25 | CD45 | Receptor-type tyrosine-protein phosphatase C | P08575 |
| 26 | CD8 | T-cell surface glycoprotein CD8 alpha / beta chain | P01732 / P10966 |
| 27 | CDC42 | Cell division control protein 42 homolog | P60953 |
| 28 | CERAMIDE | Lipid molecules involved in lipid signalling | |
| 29 | CONA | Concavallin A | CHEMBL3215495 |
| 30 | CRE | cAMP responsive element | |
| 31 | CREB1 | Cyclic AMP-responsive element-binding protein 1 | P16220 |
| 32 | CSK | Tyrosine-protein kinase CSK | P41240 |
| 33 | CYC1 | Cytochrome c1, heme protein, mitochondrial | P08574 |
| 34 | DAG1 | Diacylglycerol | |
| 35 | DGK | Diacylglycerol kinase family | |
| 36 | DMF | Dimethyl Fumarate | CHEMBL2107333 |
| 37 | EGCG | (-)-Epigallocatechin Gallate | CHEMBL297453 |
| 38 | EGF | Pro-epidermal growth factor | P01133 |
| 39 | EGFR | Epidermal growth factor receptor | P00533 |
| 40 | FAK1 | Focal adhesion kinase 1 | Q05397 |
| 41 | FCGR | Immunoglobulin gamma Fc receptor family | |
| 42 | FOS | Proto-oncogene c-Fos | P01100 |
| 43 | FOXO1 | Forkhead box protein O1 | Q12778 |
| 44 | FOXO3 | Forkhead box protein O3 | O43524 |
| 45 | FYN | Tyrosine-protein kinase Fyn | P06241 |
| 46 | GAB2 | GRB2-associated-binding protein 2 | Q9UQC2 |
| 47 | GADS | GRB2-related adapter protein 2 | O75791 |
| 48 | GAP | Ras GTPase-activating protein 1 | P20936 |
| 49 | GILENYA | Gilenya, Fingolimod | CHEMBL314854 |
| 50 | GP130 | Interleukin-6 receptor subunit beta | P40189 |
| 51 | GRB2 | Growth factor receptor-bound protein 2 | P62993 |
| 52 | GRP1 | RAS guanyl-releasing protein 1 | O95267 |
| 53 | GSK3A | Glycogen synthase kinase-3 alpha | P49840 |
| 54 | GSK3B | Glycogen synthase kinase-3 beta | P49841 |
| 55 | H2O2 | Hydrogen Peroxide | CHEMBL71595 |
| 56 | HSPB1 | Heat shock protein beta-1 | P04792 |
| 57 | ICAM1 | Intercellular adhesion molecule 1 | P05362 |
| 58 | IFNG | Interferon gamma | P01579 |
| 59 | IKBA | NF-kappa-B inhibitor alpha | P25963 |
| 60 | IKKB | Inhibitor of nuclear factor kappa-B kinase subunit beta | O14920 |
| 61 | IL1A | Interleukin-1 alpha | P01583 |
| 62 | IL1R1 | Interleukin-1 receptor type 1 | P14778 |
| 63 | IL2 | Interleukin-2 | P60568 |
| 64 | IL4 | Interleukin-4 | P05112 |
| 65 | IL6 | Interleukin-6 | P05231 |
| 66 | INS | Insulin | P01308 |
| 67 | INSR | Insulin receptor | P06213 |
| 68 | INTEGRIN | Integrin protein family | |
| 69 | IP3 | Ionositol 1,4,5-Triphosphate | CHEMBL279107 |
| 70 | IRAK4 | Interleukin-1 receptor-associated kinase 4 | Q9NWZ3 |
| 71 | IRAKM | Interleukin-1 receptor-associated kinase 3 | Q9Y616 |
| 72 | IRF1 | Interferon regulatory factor 1 | P10914 |
| 73 | ITK | Tyrosine-protein kinase ITK/TSK | Q08881 |
| 74 | JAK1 | Tyrosine-protein kinase JAK1 | P23458 |

| | | | |
|---|---|---|---|
| 75 | JAK3 | Tyrosine-protein kinase JAK3 | P52333 |
| 76 | JNK | Mitogen-activated protein kinase 8 / 9 / 10 | P45983/P45984/P53779 |
| 77 | JUN | Transcription factor AP-1 | P05412 |
| 78 | KCC2B | Calcium/calmodulin-dependent protein kinase type II subunit beta | Q13554 |
| 79 | KCC4 | Calcium/calmodulin-dependent protein kinase type IV | Q16566 |
| 80 | KEAP1 | Kelch-like ECH-associated protein 1 | Q14145 |
| 81 | KPCZ | Protein kinase C zeta type | Q05513 |
| 82 | LAT | Linker for activation of T-cells family member 1 | O43561 |
| 83 | LCK | Tyrosine-protein kinase Lck | P06239 |
| 84 | LPS | Lipopolysaccharides | |
| 85 | MALT1 | Mucosa-associated lymphoid tissue lymphoma translocation protein 1 | Q9UDY8 |
| 86 | MAP2K3 | Dual specificity mitogen-activated protein kinase kinase 3 | P46734 |
| 87 | MAP2K4 | Dual specificity mitogen-activated protein kinase kinase 4 | P45985 |
| 88 | MAP2K6 | Dual specificity mitogen-activated protein kinase kinase 6 | P52564 |
| 89 | MAP3K1 | Mitogen-activated protein kinase kinase kinase 1 | Q13233 |
| 90 | MAP3K11 | Mitogen-activated protein kinase kinase kinase 11 | Q16584 |
| 91 | MAP4K1 | Mitogen-activated protein kinase kinase kinase kinase 1 | Q92918 |
| 92 | MAPK8 | Mitogen-activated protein kinase 8 | P45983 |
| 93 | MK12 | Mitogen-activated protein kinase 12 | P53778 |
| 94 | MK03 | Mitogen-activated protein kinase 3 | P27361 |
| 95 | MP2K1 | Dual specificity mitogen-activated protein kinase kinase 1 | Q02750 |
| 96 | MRP2 | Canalicular multispecific organic anion transporter 1 | Q92887 |
| 97 | MYD88 | Myeloid differentiation primary response protein MyD88 | Q99836 |
| 98 | NACL | Sodium chloride | CHEMBL1200574 |
| 99 | NEMO | NF-kappa-B essential modulator | Q9Y6K9 |
| 100 | NF2L2 | Nuclear factor erythroid 2-related factor 2 | Q16236 |
| 101 | NFAT | Nuclear factor of activated T-cells, C1 / C2 / C3 / C4 | O95644/Q13469/Q12968/Q14934 |
| 102 | NQO1 | NAD(P)H dehydrogenase [quinone] 1 | P15559 |
| 103 | NTRK1 | High affinity nerve growth factor receptor | P04629 |
| 104 | P21 | Cyclin-dependent kinase inhibitor 1 | P38936 |
| 105 | P53 | Cellular tumor antigen p53 | P04637 |
| 106 | PAG | Phosphoprotein associated with glycosphingolipid-enriched microdomains 1 | Q9NWQ8 |
| 107 | PAR4 | PRKC apoptosis WT1 regulator protein | Q96IZ0 |
| 108 | PDPK1 | 3-phosphoinositide-dependent protein kinase 1 | O15530 |
| 109 | PGFB | Platelet-derived growth factor subunit B | P01127 |
| 110 | PGFRB | Platelet-derived growth factor receptor beta | P09619 |
| 111 | PI3K | Phosphoinositide 3-kinase family | |
| 112 | PIP3 | Phosphatidylinositol (3,4,5) - trisphosphate | CHEMBL1685065 |
| 113 | PKC | Protein kinase C family | |
| 114 | PLCB1 | 1-phosphatidylinositol 4,5-bisphosphate phosphodiesterase beta-1 | Q9NQ66 |
| 115 | PLCG1 | 1-phosphatidylinositol 4,5-bisphosphate phosphodiesterase gamma-1 | P19174 |
| 116 | POLYIC | Polyinosinic:polycytidylic acid | |
| 117 | PRDM1 | PR domain zinc finger protein 1 | O75626 |
| 118 | PSA6 | Cyclin-dependent kinase inhibitor 1B | P46527 |
| 119 | PTEN | Phosphatidylinositol 3,4,5-trisphosphate 3-phosphatase and dual-specificity protein phosphatase PTEN | P60484 |
| 120 | PTN11 | Tyrosine-protein phosphatase non - receptor type 11 | Q06124 |
| 121 | PUMA | Bcl-2-binding component 3 | Q9BXH1/Q96PG8 |
| 122 | RAC1 | Ras-related C3 botulinum toxin substrate 1 | P63000 |
| 123 | RAF1 | RAF proto-oncogene serine/threonine-protein kinase | P04049 |
| 124 | RAS | GTPase Kras / Hras / NRas | P01116/P01112/P01111 |
| 125 | REBIF | Interferon beta-1a | CHEMBL1201562 |
| 126 | RPS6KA1 | Ribosomal protein S6 kinase alpha-1 | Q15418 |
| 127 | RPS6KA5 | Ribosomal protein S6 kinase alpha-5 | O75582 |
| 128 | RPS6KB1 | Ribosomal protein S6 kinase beta-1 | P23443 |
| 129 | RS6 | 40S ribosomal protein S6 | P62753 |
| 130 | S1P1 | Sphingosine-1-phosphate | CHEMBL78494 |
| 131 | S1PR1 | Sphingosine 1-phosphate receptor 1 | P21453 |
| 132 | SGK | Serine/threonine-protein kinase Sgk1/2/3 | O00141/Q9HBY8/Q96BR1 |
| 133 | SH3BP2 | SH3 domain-binding protein 2 | P78314 |
| 134 | SHC | SHC-transforming protein 1 | P29353 |
| 135 | SHIP | Phosphatidylinositol 3,4,5-trisphosphate 5-phosphatase 1 | Q92835 |
| 136 | SHP1 | Protein-tyrosine phosphatase | P29350 |
| 137 | SLP76 | Lymphocyte cytosolic protein 2 | Q13094 |
| 138 | SOCS1 | Suppressor of cytokine signaling 1 | O15524 |
| 139 | SOCS3 | Suppressor of cytokine signaling 3 | O14543 |
| 140 | SOS | Son of sevenless homolog 1 | Q07889 |
| 141 | SRE | Serum response element | |
| 142 | STAT1 | Signal transducer and activator of transcription 1 | P42224 |
| 143 | STAT2 | Signal transducer and activator of transcription 2 | P52630 |
| 144 | STAT3 | Signal transducer and activator of transcription 3 | P40763 |
| 145 | STAT5 | Signal transducer and activator of transcription 5A / 5B | P42229/P51692 |
| 146 | STAT6 | Signal transducer and activator of transcription 6 | P42226 |
| 147 | SYK | Tyrosine-protein kinase SYK | P43405 |
| 148 | TAK1 | Mitogen-activated protein kinase kinase kinase 7 | O43318 |
| 149 | TCR | T-cell Receptor | |

| | | | |
|---|---|---|---|
| 150 | TCRP | T-cell Receptor phosphorylated | |
| 151 | TERIFLUNOMIDE | Teriflunomide, Aubagio | CHEMBL973 |
| 152 | TF65 | Transcription factor p65 | Q04206 |
| 153 | TLR3 | Toll-like receptor 3 | O15455 |
| 154 | TLR4 | Toll-like receptor 4 | O00206 |
| 155 | TNFA | Tumor necrosis factor | P01375 |
| 156 | TOLLIP | Toll-interacting protein | Q9H0E2 |
| 157 | TRADD | Tumor necrosis factor receptor type 1-associated DEATH domain protein | Q15628 |
| 158 | TRAF2 | TNF receptor-associated factor 2 | Q12933 |
| 159 | TRAF6 | TNF receptor-associated factor 6 | Q9Y4K3 |
| 160 | TXK | Tyrosine-protein kinase TXK | P42681 |
| 161 | VAV | Proto-oncogene vav | P15498 |
| 162 | VAV3 | Guanine nucleotide exchange factor VAV3 | Q9UKW4 |
| 163 | VDR | Vitamin D3 receptor | P11473 |
| 164 | VITD3 | Vitamin D 3, Cholecalciferol | CHEMBL1042 |
| 165 | WNK1 | Serine/threonine-protein kinase WNK1 | Q9H4A3 |
| 166 | X | Non-identified kinase involved in CD28- mediated signalling, see Saez-Rodriguez2007 | |
| 167 | ZAP70 | Tyrosine-protein kinase ZAP-70 | P43403 |

**Table S3. Selection of phosphoproteins based in their involvement in pathways associated with MS or MS drugs and assays performance.** Phosphoproteins for the xMAP assays were selected based in a good signal to noise ratio in order to maximize network coverage and model identifiability.

| Protein | Phosphosite | Pathway relevance | Consensus | Multiplex-ability | SNR | Selection |
|---|---|---|---|---|---|---|
| AKT1 | S473 | HIGH | LOW | HIGH | LOW | PASS |
| CREB1 | S133 | HIGH | MEDIUM | HIGH | LOW | PASS |
| FAK1 | Y397 | MEDIUM | | LOW | LOW | PASS |
| GSK3A | S21 | MEDIUM | HIGH | MEDIUM | | PASS |
| HSPB1 | S78/S82 | MEDIUM | HIGH | | HIGH | PASS |
| IKBA | S32 | HIGH | HIGH | HIGH | LOW | PASS |
| JUN | S63 | HIGH | | MEDIUM | | PASS |
| MK03 | T202/Y204 | HIGH | MEDIUM | HIGH | MEDIUM | PASS |
| MK12 | T180/Y182 | HIGH | MEDIUM | HIGH | | PASS |
| MP2K1 | S217/S221 | HIGH | | MEDIUM | HIGH | PASS |
| PTN11 | Y542 | MEDIUM | | LOW | HIGH | PASS |
| STAT1 | Y701 | HIGH | HIGH | HIGH | | PASS |
| STAT3 | Y705 | HIGH | LOW | HIGH | | PASS |
| STAT5 | S694 | HIGH | | LOW | | PASS |
| STAT6 | Y641 | HIGH | | MEDIUM | | PASS |
| TF65 | S536 | HIGH | | | HIGH | PASS |
| WNK1 | T60 | HIGH | | HIGH | MEDIUM | PASS |
| EGFR | Y1068 | HIGH | | HIGH | HIGH | FAIL |
| GSK3B | S9 | MEDIUM | | HIGH | LOW | FAIL |
| KS6A1 | S380 | LOW | MEDIUM | HIGH | HIGH | FAIL |
| KS6B1 | T389 | LOW | HIGH | HIGH | LOW | FAIL |
| LAT | Y191 | MEDIUM | | LOW | | FAIL |
| LCK | Y505 | MEDIUM | | MEDIUM | | FAIL |
| MK09 | T183/Y185 | LOW | MEDIUM | HIGH | LOW | FAIL |
| MP2K6 | S207/T211 | LOW | | LOW | LOW | FAIL |
| NRF2 | S40 | LOW | | HIGH | | FAIL |
| P53 | S46 | MEDIUM | | HIGH | LOW | FAIL |
| PGFRB | Y751 | LOW | | HIGH | | FAIL |
| RS6 | S235/S236 | MEDIUM | | MEDIUM | LOW | FAIL |
| STAT2 | Y690 | HIGH | | LOW | | FAIL |
| VDR | S208 | LOW | | HIGH | | FAIL |
| ZAP70 | Y319 | MEDIUM | | MEDIUM | | FAIL |

**Table S4: List of phosphoproteins used for the in vitro multiplex assays with PBMCs**

| Uniprot ID Name | Entrez-gene identifier | HGNC symbol | Uniprot Recommended Name | Uniprot Alternative Name | Gene Names | Antibody company catalog | Pathway | Biological role and Association with MS |
|---|---|---|---|---|---|---|---|---|
| **AKT1** | 207 | AKT1 | RAC-alpha serine/threonine-protein kinase | Protein kinase B | AKT1, PKB, RAC | PAO P-AKT1-A01 | PI3K/AKT/mTOR | key mediator of PI3K and mTOR signaling pathways, all involved in cell survival |
| **CREB1** | 1385 | CREB1 | Cyclic AMP-responsive element-binding protein 1 | - | CREB1 | PAO: P-CREB1-A01 | AP-1/MAPK | Participates in PI3K, MAPKinase pathways, promoting cell survival, neuronal activity, synaptic plasticity and expression of HLA molecules |
| **FAK1** | 5747 | PTK2 | Focal adhesion kinase 1 | Protein phosphatase 1 regulatory subunit 71 | PTK2, FAK, FAK1 | PAO: P-FAK1-A01 | Integrin/src | key signaling mediator of integrin receptors and axon guidance as well as growth factor receptors (PI3K pathway) |
| **GSK3A** | 2931 | GSK3A | Glycogen synthase kinase-3 alpha | Serine/threonine-protein kinase GSK3A | GSK3A | PAO: P-GSK3A-A01 | PI3K/AKT/mTOR | chemokine signaling pathway, being downstream AKT |
| **HSPB1** | 3315 | HSPB1 | Heat shock protein beta-1 | 28 kDa heat shock protein | HSPB1, HSP27, HSP28 | PAO: P-HSPB1-A01 | p38/MAPK | Part of MAPKinase pathway, downstream p38, and promotes actin reorganization supporing cell migratio. Marked elevation in HSP27 levels during the relapse phase of MS. |
| **IKBA** | 4792 | NFKBIA | NF-kappa-B inhibitor alpha | I-kappa-B-alpha | NFKBIA, IKBA, MAD3, NFKBI | PAO: P-IKBA-A01 | NFkB | NFkB signaling |

| | | | | | | | | |
|---|---|---|---|---|---|---|---|---|
| **JUN** | 3725 | JUN | Transcription factor AP-1 | Proto-oncogene c-Jun | JUN | PAO: P-JUN-A01 | JNK/MAPK | JUN is part of the immediate early gene responses, being downstream JNK and mediating apoptosis |
| **MK03** | 5595 | MAPK3 | Mitogen-activated protein kinase 3 | ERK1 | MAPK3, ERK1, PRKM3 | PAO: P-MK03-A01 | AP-1/MAPK | Member of the MAPKinase pathway, involved in proliferation, differentiation, and cell cycle progression |
| **MK12** | 6300 | MAPK12 | Mitogen-activated protein kinase 12 | MAP kinase p38 gamma | MAPK12, ERK6, SAPK3 | PAO: P-MK12-A01 | AP-1/MAPK | Member of the MAPKinase pathway, involved in proliferation, differentiation, and cell cycle progression |
| **MP2K1** | 5604 | MAP2K1 | Dual specificity mitogen-activated protein kinase kinase 1 | MEK1 | MAP2K1, MEK1, PRKMK1 | PAO: P-MP2K1-A01 | AP-1/MAPK | Mediates signaling of PDGFR pathway, promoting cell survival |
| **PTPN11** | 5781 | PTPN11 | Tyrosine-protein phosphatase non-receptor type 11 | SHP2 | PTPN11, PTP2C, SHPTP2 | PAO: P-PTN11-A01 | AP-1/MAPK | Member of the MAPKinase pathway, involved in proliferation, differentiation, cell cycle progression, cell transmigration, NK cytotxicity and axonal guidance |
| **STAT5A** | 6776 | STAT5A | Signal transducer and activator of transcription 5A | - | STAT5A, STAT5 | PAO: P-STAT5-A01 | JAK/STAT | Mediates IL-2, chemokines, NGR3-ErbB signaling and promoting cell survival. Upon IL-7 stimulation, MS patients experience stronger STAT5 activation in CD8-EM compared with HC. |
| **STAT1** | 6772 | STAT1 | Signal transducer and activator of transcription 1-alpha/beta | Transcription factor ISGF-3 components p91/p84 | STAT1 | PAO: P-STAT1-A01 | JAK/STAT | Mediates interferon gamma and beta signaling. Macrophages from MS patients displayed enhanced STAT1, STAT6 and NF-kB activity. |

| | | | | | | | | |
|---|---|---|---|---|---|---|---|---|
| **STAT3** | 6774 | STAT3 | Signal transducer and activator of transcription 3 | Acute-phase response factor | STAT3, APRF | PAO: P-STAT3-A01 | JAK/STAT | Mediates IL-6, IL-10, neurocytokines (LIF) signaling. STAT3 is required for IL-17 production by Th17. |
| **STAT6** | 6778 | STAT6 | Signal transducer and activator of transcription 6 | IL-4 Stat | STAT6 | PAO: P-STAT6-A01 | JAK/STAT | JAK-STAT pathway mediates Il-4 signaling. PBMCs from MS patients have significantly elevated constitutive phosphorylation of STAT6 compared to PBMCs from normal subjects. |
| **TF65** | 5970 | RELA | Transcription factor p65 | Nuclear factor NF-kappa-B p65 subunit | RELA, NFKB3 | PAO: P-TF65-A01 | NFkB | Key member of NfKB pathway, meadiating inflammatory signals and cell survival. Macrophages from MS patients displayed enhanced STAT6, STAT1 and NF-kB activity. |
| **WNK1** | 65125 | WNK1 | Serine/threonine-protein kinase WNK1 | Erythrocyte 65 kDa protein | WNK1, HSN2, KDP, KIAA0344, PRKWNK1 | PAO: P-WNK1-A01 | ERK5/MAPK | EGF pathway and participate in the regulation of ion homeostasis and iron uptake |

**Table S5: Stimuli used for the in vitro multiplex assays with PBMCs**

| Stimulus | Full Name | pathway | cat number | Vendor | stock conc | units | target conc | units |
|---|---|---|---|---|---|---|---|---|
| antiCD3 | Anti-Human CD3 | TCR | 16-0037-85 | eBioscience | 1 | mg/ml | 5000 | ng/ml |
| BDNF | Brain-Derived Neurotrophic Factor human | TrkB/p75 | B3795-5ug | Sigma | 0.1 | mg/ml | 100 | ng/ml |
| conA | Concanavalin A | TCR | C5275-5MG | Sigma | 5 | mg/ml | 2500 | ng/ml |
| DMF | Dimethyl fumarate | Nrf2 | 242926 | Sigma | 5 | mg/ml | 10000 | ng/ml |
| EGCG | Epigallocatechin-3-gallate | anti-oxidative stress | sc-200802 | Santa Cruz | 10 | mg/ml | 45.84 | ug/ml |
| FTY | Fingolimod | S1PR | | Novartis | 1 | mg/ml | 3000 | ng/ml |
| H2O2 | Hydrogen peroxide | oxidative stress | H3410 | Sigma | 330 | mg/ml | 17005 | ng/ml |
| IFNB1a | Interferon beta 1a | Type I IFNR | 101322 | Merck | 0.088 | mg/ml | 50 | ng/ml |
| IFNG | Interferon gamma | Type II IFNR | I3265 | Sigma | 0.1 | mg/ml | 100 | ng/ml |
| IL1A | Interleukin-1 alpha | IL1R pathway | 200-01A | PeproTech | 0.1 | mg/ml | 50 | ng/ml |
| IL6 | Interleukin-6 | IL6R | 200-06 | PeproTech | 0.1 | mg/ml | 100 | ng/ml |

| | | | | | | | | |
|---|---|---|---|---|---|---|---|---|
| INS | Insulin | InsulinR | I9278 | Sigma | 1.722 | mg/ml | 1722 | ng/ml |
| LPS | Lipopolysaccharide | TLR4 | L4391 | Sigma | 1 | mg/ml | 10000 | ng/ml |
| NaCl | Sodium Chloride | PI3K | S5886 | Sigma | 11.7 | mg/ml | 2.34 | mg/ml |
| PolyIC | Polyinosinic-Polycytidylic acid | TLR3 | p0913-10mg | Sigma | 10 | mg/ml | 10000 | ng/ml |
| S1P | Sphingosine 1-phosphate | S1P | S9666 | Sigma | 0.125 | mg/ml | 100 | ng/ml |
| Teriflu | Teriflunomide | pyrimidin synthesis NFkB inhibitor | A77 1726 | Calbiochem | 10 | mg/ml | 13510 | ng/ml |
| TNFA | Tumor necrosis factor alpha | TNF | 300-01A | PeproTech | 0.1 | mg/ml | 100 | ng/ml |
| vitD3 | vitamin D3 | B-catenin | C9756 | Sigma | 1 | mg/ml | 500 | ng/ml |
| BN201 | Neuroprotective peptoid IGF-1 NA Bionure Farma | SGK | Under preparation | Bionure | 1 | mg/ml | 5000 | ng/ml |

**Table S6. Demographics and clinical variables of MS patients and controls.**

| | MS<br>n=195 | HC<br>n=60 |
|---|---|---|
| **Sex (M/F)** | 66/129 | 21/39 |
| **Age** | 43.1±11.3 | 39.9±8.5* |
| **Disease duration (months)** | 104.9+93.2 | -- |
| **Age at onset** | 34.5+10.3 | -- |
| **Disease subtype** | | |
| **CIS** | 24 | -- |
| **RRMS** | 129 | -- |
| **SPMS** | 6 | -- |
| **PPMS** | 36 | -- |
| **EDSS** | 2 (0-6.0) | -- |
| **DMD** | | |
| **IFNb** | 37 | -- |
| **GA** | 18 | -- |
| **NTZ** | 22 | -- |
| **FTY** | 20 | -- |
| **Untreated** | 98 | -- |

**Table S7. List of co-druggable interactions and co-druggability score.** By our definition, co-druggable interactions must have score smaller or equal to zero.

| No | ECGC | | FTY | | GA | | IFNbeta | | NTZ | |
|---|---|---|---|---|---|---|---|---|---|---|
| 1 | !LCK=PI3K | -0.41 | !STAT3=GP130 | -0.52 | CONA=GRB2 | -0.36 | !CBL=ZAP70 | -0.19 | !ANTICD3=PAG | -0.06 |
| 2 | !LCK=STAT3 | -0.33 | AKT1=GSK3A | -0.19 | DAG1=RAS | -0.23 | BDNF=GRB2 | 0.00 | ANTICD3=WNK1 | 0.00 |
| 3 | AKT1=GSK3A | -0.34 | ANTICD3=TCRP | -0.17 | IL1A=IL1R1 | -0.14 | CONA=VAV | 0.00 | BDNF=GRB2 | -0.05 |
| 4 | BDNF=GRB2 | -0.23 | BDNF=GRB2 | -0.14 | IL1R1=TAK1 | -0.16 | GP130=JAK1 | -0.11 | CONA=STAT1 | -0.06 |
| 5 | CONA=GRB2 | -0.33 | IL1A=IL1R1 | -0.11 | MAP2K4=MK12 | -0.16 | JAK1=STAT1 | -0.06 | GP130=JAK1 | -0.10 |
| 6 | CONA=VAV | -0.42 | IL1R1=TAK1 | -0.12 | MAP3K1=MAP2K4 | -0.19 | LPS=TAK1 | -0.14 | INS=PI3K | 0.00 |
| 7 | GRB2=RAS | -0.28 | IL6=GP130 | -0.43 | NACL=MK12 | -0.12 | MAP3K1=MAP2K4 | -0.08 | MAP2K4=MK12 | -0.15 |
| 8 | JAK1=STAT3 | 0.00 | MK12=HSPB1 | -0.28 | PAG=LCK | -0.22 | MAP3K1=MK12 | -0.12 | MAP3K1=MAP2K4 | -0.08 |
| 9 | PI3K=PIP3 | -0.36 | NACL=MK12 | 0.00 | PIP3=PLCG1 | -0.26 | NACL=MK12 | -0.16 | NACL=MK12 | -0.06 |
| 10 | PIP3=AKT1 | -0.31 | RAC1=MAP3K1 | -0.19 | PLCG1=DAG1 | -0.26 | PAG=LCK | -0.13 | REBIF=JAK1 | -0.05 |
| 11 | RAS=MAP3K1 | -0.57 | REBIF=JAK1 | -0.09 | RAF1=SGK | -0.26 | SLP76=AKT1 | -0.14 | SLP76=AKT1 | -0.10 |
| 12 | TAK1=MAP2K4 | -0.36 | TCRP=ZAP70 | -0.28 | SLP76=AKT1 | -0.28 | ZAP70=GRB2 | 0.00 | TAK1=MK12 | -0.15 |
| 13 | VAV=RAC1 | -0.37 | | | ZAP70=SLP76 | -0.26 | ZAP70=SLP76 | 0.00 | | |

**Table S8. Predicted targets for combination therapy.** The drug target is the parent of the interaction. The stimulus is the one that triggers the interaction between both kinases. Co-druggability score is always smaller or equal to zero in order to be selected as co-druggable interactions. Boolean network activity represents the active (1) and inactive (0) reactions for each subgroup (**Figure 5**). Group network activity denotes the mean signaling activity for each drug-treated subgroup (**Supplementary Figure S3**).

| | Reaction | Stimulus | Readout | drugScore | Boolean network activity | Group network activity |
|---|---|---|---|---|---|---|
| | | | **EGCG** | | | |
| 1 | AKT1=GSK3A | INS | GSK3A | -0.34 | 0 | 0.00 |
| 2 | BDNF=GRB2 | BDNF | MK12 | -0.23 | 1 | 1.00 |
| 3 | BDNF=GRB2 | BDNF | MP2K1 | -0.23 | 1 | 1.00 |
| 4 | BDNF=GRB2 | BDNF | MKO3 | -0.23 | 1 | 1.00 |
| 5 | CONA=GRB2 | CONA | MK12 | -0.33 | 1 | 0.67 |
| 6 | CONA=GRB2 | CONA | MP2K1 | -0.33 | 1 | 0.67 |
| 7 | CONA=GRB2 | CONA | MKO3 | -0.33 | 1 | 0.67 |
| 8 | GRB2=RAS | BDNF | MK12 | -0.28 | 1 | 1.00 |
| 9 | GRB2=RAS | BDNF | MP2K1 | -0.28 | 1 | 1.00 |
| 10 | GRB2=RAS | BDNF | MKO3 | -0.28 | 1 | 1.00 |
| 11 | GRB2=RAS | IL1A | MK12 | -0.28 | 1 | 1.00 |
| 12 | GRB2=RAS | IL1A | MP2K1 | -0.28 | 1 | 1.00 |
| 13 | GRB2=RAS | IL1A | MKO3 | -0.28 | 1 | 1.00 |
| 14 | GRB2=RAS | LPS | MK12 | -0.28 | 1 | 1.00 |
| 15 | GRB2=RAS | LPS | MP2K1 | -0.28 | 1 | 1.00 |
| 16 | GRB2=RAS | LPS | MKO3 | -0.28 | 1 | 1.00 |
| 17 | GRB2=RAS | POLYIC | MK12 | -0.28 | 1 | 1.00 |
| 18 | GRB2=RAS | POLYIC | MP2K1 | -0.28 | 1 | 1.00 |
| 19 | GRB2=RAS | POLYIC | MKO3 | -0.28 | 1 | 1.00 |
| 20 | GRB2=RAS | CONA | MK12 | -0.28 | 1 | 1.00 |
| 21 | GRB2=RAS | CONA | MP2K1 | -0.28 | 1 | 1.00 |
| 22 | GRB2=RAS | CONA | MKO3 | -0.28 | 1 | 1.00 |
| 23 | GRB2=RAS | TNFA | MK12 | -0.28 | 1 | 1.00 |
| 24 | GRB2=RAS | TNFA | MP2K1 | -0.28 | 1 | 1.00 |
| 25 | GRB2=RAS | TNFA | MKO3 | -0.28 | 1 | 1.00 |
| 26 | JAK1=STAT3 | REBIF | STAT3 | 0.00 | 1 | 0.67 |
| 27 | JAK1=STAT3 | IL6 | STAT3 | 0.00 | 1 | 0.67 |
| 28 | PI3K=PIP3 | INS | AKT1 | -0.36 | 0 | 0.33 |
| 29 | PI3K=PIP3 | INS | GSK3A | -0.36 | 0 | 0.33 |
| 30 | PIP3=AKT1 | INS | AKT1 | -0.31 | 0 | 0.00 |
| 31 | PIP3=AKT1 | INS | GSK3A | -0.31 | 0 | 0.00 |
| 32 | RAS=MAP3K1 | BDNF | MK12 | -0.57 | 1 | 1.00 |
| 33 | RAS=MAP3K1 | BDNF | MP2K1 | -0.57 | 1 | 1.00 |
| 34 | RAS=MAP3K1 | BDNF | MKO3 | -0.57 | 1 | 1.00 |
| 35 | RAS=MAP3K1 | IL1A | MK12 | -0.57 | 1 | 1.00 |
| 36 | RAS=MAP3K1 | IL1A | MP2K1 | -0.57 | 1 | 1.00 |

| | | | | | | |
|---|---|---|---|---|---|---|
| 36 | RAS=MAP3K1 | IL1A | MP2K1 | -0.57 | 1 | 1.00 |
| 37 | RAS=MAP3K1 | IL1A | MKO3 | -0.57 | 1 | 1.00 |
| 38 | RAS=MAP3K1 | LPS | MK12 | -0.57 | 1 | 1.00 |
| 39 | RAS=MAP3K1 | LPS | MP2K1 | -0.57 | 1 | 1.00 |
| 40 | RAS=MAP3K1 | LPS | MKO3 | -0.57 | 1 | 1.00 |
| 41 | RAS=MAP3K1 | POLYIC | MK12 | -0.57 | 1 | 1.00 |
| 42 | RAS=MAP3K1 | POLYIC | MP2K1 | -0.57 | 1 | 1.00 |
| 43 | RAS=MAP3K1 | POLYIC | MKO3 | -0.57 | 1 | 1.00 |
| 44 | RAS=MAP3K1 | CONA | MK12 | -0.57 | 1 | 1.00 |
| 45 | RAS=MAP3K1 | CONA | MP2K1 | -0.57 | 1 | 1.00 |
| 46 | RAS=MAP3K1 | CONA | MKO3 | -0.57 | 1 | 1.00 |
| 47 | RAS=MAP3K1 | TNFA | MK12 | -0.57 | 1 | 1.00 |
| 48 | RAS=MAP3K1 | TNFA | MP2K1 | -0.57 | 1 | 1.00 |
| 49 | RAS=MAP3K1 | TNFA | MKO3 | -0.57 | 1 | 1.00 |
| 50 | TAK1=MAP2K4 | IL1A | MK12 | -0.36 | 1 | 0.67 |
| 51 | TAK1=MAP2K4 | IL1A | MP2K1 | -0.36 | 1 | 0.67 |
| 52 | TAK1=MAP2K4 | IL1A | MKO3 | -0.36 | 1 | 0.67 |
| 53 | TAK1=MAP2K4 | LPS | MK12 | -0.36 | 1 | 0.67 |
| 54 | TAK1=MAP2K4 | LPS | MP2K1 | -0.36 | 1 | 0.67 |
| 55 | TAK1=MAP2K4 | LPS | MKO3 | -0.36 | 1 | 0.67 |
| 56 | TAK1=MAP2K4 | POLYIC | MK12 | -0.36 | 1 | 0.67 |
| 57 | TAK1=MAP2K4 | POLYIC | MP2K1 | -0.36 | 1 | 0.67 |
| 58 | TAK1=MAP2K4 | POLYIC | MKO3 | -0.36 | 1 | 0.67 |
| **FTY** | | | | | | |
| 1 | AKT1=GSK3A | BDNF | GSK3A | -0.19 | 0 | 0.15 |
| 2 | AKT1=GSK3A | NACL | GSK3A | -0.19 | 0 | 0.15 |
| 3 | AKT1=GSK3A | IL1A | GSK3A | -0.19 | 0 | 0.15 |
| 4 | AKT1=GSK3A | INS | GSK3A | -0.19 | 0 | 0.15 |
| 5 | AKT1=GSK3A | LPS | GSK3A | -0.19 | 0 | 0.15 |
| 6 | AKT1=GSK3A | POLYIC | GSK3A | -0.19 | 0 | 0.15 |
| 7 | AKT1=GSK3A | CONA | GSK3A | -0.19 | 0 | 0.15 |
| 8 | AKT1=GSK3A | TNFA | GSK3A | -0.19 | 0 | 0.15 |
| 9 | BDNF=GRB2 | BDNF | HSPB1 | -0.14 | 0 | 0.38 |
| 10 | BDNF=GRB2 | BDNF | AKT1 | -0.14 | 0 | 0.38 |
| 11 | BDNF=GRB2 | BDNF | MK12 | -0.14 | 0 | 0.38 |
| 12 | BDNF=GRB2 | BDNF | GSK3A | -0.14 | 0 | 0.38 |
| 13 | BDNF=GRB2 | BDNF | MP2K1 | -0.14 | 0 | 0.38 |
| 14 | BDNF=GRB2 | BDNF | MKO3 | -0.14 | 0 | 0.38 |
| 15 | BDNF=GRB2 | BDNF | STAT5 | -0.14 | 0 | 0.38 |
| 16 | IL1A=IL1R1 | IL1A | HSPB1 | -0.11 | 1 | 0.54 |
| 17 | IL1A=IL1R1 | IL1A | AKT1 | -0.11 | 1 | 0.54 |
| 18 | IL1A=IL1R1 | IL1A | MK12 | -0.11 | 1 | 0.54 |
| 19 | IL1A=IL1R1 | IL1A | GSK3A | -0.11 | 1 | 0.54 |
| 20 | IL1A=IL1R1 | IL1A | MP2K1 | -0.11 | 1 | 0.54 |
| 21 | IL1A=IL1R1 | IL1A | MKO3 | -0.11 | 1 | 0.54 |
| 22 | IL1A=IL1R1 | IL1A | STAT5 | -0.11 | 1 | 0.54 |
| 23 | IL1R1=TAK1 | IL1A | HSPB1 | -0.12 | 1 | 0.54 |
| 24 | IL1R1=TAK1 | IL1A | AKT1 | -0.12 | 1 | 0.54 |
| 25 | IL1R1=TAK1 | IL1A | MK12 | -0.12 | 1 | 0.54 |
| 26 | IL1R1=TAK1 | IL1A | GSK3A | -0.12 | 1 | 0.54 |
| 27 | IL1R1=TAK1 | IL1A | MP2K1 | -0.12 | 1 | 0.54 |
| 28 | IL1R1=TAK1 | IL1A | MKO3 | -0.12 | 1 | 0.54 |
| 29 | IL1R1=TAK1 | IL1A | STAT5 | -0.12 | 1 | 0.54 |

| | | | | | | |
|---|---|---|---|---|---|---|
| 30 | IL6=GP130 | IL6 | STAT6 | -0.43 | 0 | 0.31 |
| 31 | MK12=HSPB1 | BDNF | HSPB1 | -0.28 | 1 | 0.85 |
| 32 | MK12=HSPB1 | NACL | HSPB1 | -0.28 | 1 | 0.85 |
| 33 | MK12=HSPB1 | IL1A | HSPB1 | -0.28 | 1 | 0.85 |
| 34 | MK12=HSPB1 | INS | HSPB1 | -0.28 | 1 | 0.85 |
| 35 | MK12=HSPB1 | LPS | HSPB1 | -0.28 | 1 | 0.85 |
| 36 | MK12=HSPB1 | POLYIC | HSPB1 | -0.28 | 1 | 0.85 |
| 37 | MK12=HSPB1 | CONA | HSPB1 | -0.28 | 1 | 0.85 |
| 38 | MK12=HSPB1 | TNFA | HSPB1 | -0.28 | 1 | 0.85 |
| 39 | NACL=MK12 | NACL | HSPB1 | 0.00 | 0 | 0.31 |
| 40 | NACL=MK12 | NACL | AKT1 | 0.00 | 0 | 0.31 |
| 41 | NACL=MK12 | NACL | MK12 | 0.00 | 0 | 0.31 |
| 42 | NACL=MK12 | NACL | GSK3A | 0.00 | 0 | 0.31 |
| 43 | NACL=MK12 | NACL | MP2K1 | 0.00 | 0 | 0.31 |
| 44 | NACL=MK12 | NACL | MKO3 | 0.00 | 0 | 0.31 |
| 45 | NACL=MK12 | NACL | STAT5 | 0.00 | 0 | 0.31 |
| 46 | RAC1=MAP3K1 | CONA | HSPB1 | -0.19 | 1 | 0.46 |
| 47 | RAC1=MAP3K1 | CONA | AKT1 | -0.19 | 1 | 0.46 |
| 48 | RAC1=MAP3K1 | CONA | MK12 | -0.19 | 1 | 0.46 |
| 49 | RAC1=MAP3K1 | CONA | GSK3A | -0.19 | 1 | 0.46 |
| 50 | RAC1=MAP3K1 | CONA | MP2K1 | -0.19 | 1 | 0.46 |
| 51 | RAC1=MAP3K1 | CONA | MKO3 | -0.19 | 1 | 0.46 |
| 52 | RAC1=MAP3K1 | CONA | STAT5 | -0.19 | 1 | 0.46 |
| 53 | REBIF=JAK1 | REBIF | STAT6 | -0.09 | 1 | 0.54 |
| **GA** | | | | | | |
| 1 | CONA=GRB2 | CONA | MP2K1 | -0.36 | 1 | 0.70 |
| 2 | CONA=GRB2 | CONA | MKO3 | -0.36 | 1 | 0.70 |
| 3 | DAG1=RAS | INS | MP2K1 | -0.23 | 0 | 0.30 |
| 4 | DAG1=RAS | INS | MKO3 | -0.23 | 0 | 0.30 |
| 5 | IL1A=IL1R1 | IL1A | MK12 | -0.14 | 1 | 0.50 |
| 6 | IL1A=IL1R1 | IL1A | MKO3 | -0.14 | 1 | 0.50 |
| 7 | IL1R1=TAK1 | IL1A | MK12 | -0.16 | 1 | 0.50 |
| 8 | IL1R1=TAK1 | IL1A | MKO3 | -0.16 | 1 | 0.50 |
| 9 | MAP2K4=MK12 | TNFA | MK12 | -0.16 | 0 | 0.20 |
| 10 | MAP2K4=MK12 | TNFA | MKO3 | -0.16 | 0 | 0.20 |
| 11 | MAP3K1=MAP2K4 | TNFA | MK12 | -0.19 | 0 | 0.10 |
| 12 | MAP3K1=MAP2K4 | TNFA | MKO3 | -0.19 | 0 | 0.10 |
| 13 | NACL=MK12 | NACL | MK12 | -0.12 | 0 | 0.20 |
| 14 | NACL=MK12 | NACL | MKO3 | -0.12 | 0 | 0.20 |
| 15 | PIP3=PLCG1 | INS | MP2K1 | -0.26 | 0 | 0.10 |
| 16 | PIP3=PLCG1 | INS | MKO3 | -0.26 | 0 | 0.10 |
| 17 | PLCG1=DAG1 | INS | MP2K1 | -0.26 | 0 | 0.30 |
| 18 | PLCG1=DAG1 | INS | MKO3 | -0.26 | 0 | 0.30 |
| 19 | RAF1=SGK | ANTICD3 | MKO3 | -0.26 | 1 | 0.50 |
| 20 | RAF1=SGK | BDNF | MKO3 | -0.26 | 1 | 0.50 |
| 21 | RAF1=SGK | INS | MKO3 | -0.26 | 1 | 0.50 |
| 22 | RAF1=SGK | CONA | MKO3 | -0.26 | 1 | 0.50 |
| 23 | SLP76=AKT1 | ANTICD3 | AKT1 | -0.28 | 1 | 0.60 |
| 24 | SLP76=AKT1 | ANTICD3 | STAT1 | -0.28 | 1 | 0.60 |
| 25 | SLP76=AKT1 | ANTICD3 | GSK3A | -0.28 | 1 | 0.60 |
| 26 | ZAP70=SLP76 | ANTICD3 | AKT1 | -0.26 | 1 | 0.60 |
| 27 | ZAP70=SLP76 | ANTICD3 | STAT1 | -0.26 | 1 | 0.60 |
| 28 | ZAP70=SLP76 | ANTICD3 | GSK3A | -0.26 | 1 | 0.60 |

| IFNb | | | | | | |
|---|---|---|---|---|---|---|
| 1 | !CBL=ZAP70 | EGCG | HSPB1 | -0.19 | 1 | 0.50 |
| 2 | !CBL=ZAP70 | EGCG | AKT1 | -0.19 | 1 | 0.50 |
| 3 | !CBL=ZAP70 | EGCG | MK12 | -0.19 | 1 | 0.50 |
| 4 | !CBL=ZAP70 | EGCG | GSK3A | -0.19 | 1 | 0.50 |
| 5 | !CBL=ZAP70 | EGCG | MP2K1 | -0.19 | 1 | 0.50 |
| 6 | !CBL=ZAP70 | EGCG | MKO3 | -0.19 | 1 | 0.50 |
| 7 | BDNF=GRB2 | BDNF | HSPB1 | 0.00 | 1 | 0.50 |
| 8 | BDNF=GRB2 | BDNF | MK12 | 0.00 | 1 | 0.50 |
| 9 | BDNF=GRB2 | BDNF | MP2K1 | 0.00 | 1 | 0.50 |
| 10 | BDNF=GRB2 | BDNF | MKO3 | 0.00 | 1 | 0.50 |
| 11 | GP130=JAK1 | IL6 | STAT1 | -0.11 | 1 | 0.96 |
| 12 | GP130=JAK1 | IL6 | STAT3 | -0.11 | 1 | 0.96 |
| 13 | JAK1=STAT1 | REBIF | STAT1 | -0.06 | 1 | 0.67 |
| 14 | JAK1=STAT1 | IFNG | STAT1 | -0.06 | 1 | 0.67 |
| 15 | JAK1=STAT1 | IL6 | STAT1 | -0.06 | 1 | 0.67 |
| 16 | LPS=TAK1 | LPS | HSPB1 | -0.14 | 1 | 0.96 |
| 17 | LPS=TAK1 | LPS | MK12 | -0.14 | 1 | 0.96 |
| 18 | LPS=TAK1 | LPS | MKO3 | -0.14 | 1 | 0.96 |
| 19 | MAP3K1=MAP2K4 | ANTICD3 | HSPB1 | -0.08 | 0 | 0.21 |
| 20 | MAP3K1=MAP2K4 | ANTICD3 | MK12 | -0.08 | 0 | 0.21 |
| 21 | MAP3K1=MAP2K4 | ANTICD3 | MKO3 | -0.08 | 0 | 0.21 |
| 22 | MAP3K1=MAP2K4 | BDNF | HSPB1 | -0.08 | 0 | 0.21 |
| 23 | MAP3K1=MAP2K4 | BDNF | MK12 | -0.08 | 0 | 0.21 |
| 24 | MAP3K1=MAP2K4 | BDNF | MKO3 | -0.08 | 0 | 0.21 |
| 25 | MAP3K1=MAP2K4 | INS | HSPB1 | -0.08 | 0 | 0.21 |
| 26 | MAP3K1=MAP2K4 | INS | MK12 | -0.08 | 0 | 0.21 |
| 27 | MAP3K1=MAP2K4 | INS | MKO3 | -0.08 | 0 | 0.21 |
| 28 | MAP3K1=MAP2K4 | EGCG | HSPB1 | -0.08 | 0 | 0.21 |
| 29 | MAP3K1=MAP2K4 | EGCG | MK12 | -0.08 | 0 | 0.21 |
| 30 | MAP3K1=MAP2K4 | EGCG | MKO3 | -0.08 | 0 | 0.21 |
| 31 | MAP3K1=MAP2K4 | CONA | HSPB1 | -0.08 | 0 | 0.21 |
| 32 | MAP3K1=MAP2K4 | CONA | MK12 | -0.08 | 0 | 0.21 |
| 33 | MAP3K1=MAP2K4 | CONA | MKO3 | -0.08 | 0 | 0.21 |
| 34 | MAP3K1=MAP2K4 | TNFA | HSPB1 | -0.08 | 0 | 0.21 |
| 35 | MAP3K1=MAP2K4 | TNFA | MK12 | -0.08 | 0 | 0.21 |
| 36 | MAP3K1=MAP2K4 | TNFA | MKO3 | -0.08 | 0 | 0.21 |
| 37 | MAP3K1=MK12 | ANTICD3 | HSPB1 | -0.12 | 1 | 0.79 |
| 38 | MAP3K1=MK12 | ANTICD3 | MK12 | -0.12 | 1 | 0.79 |
| 39 | MAP3K1=MK12 | ANTICD3 | MKO3 | -0.12 | 1 | 0.79 |
| 40 | MAP3K1=MK12 | BDNF | HSPB1 | -0.12 | 1 | 0.79 |
| 41 | MAP3K1=MK12 | BDNF | MK12 | -0.12 | 1 | 0.79 |
| 42 | MAP3K1=MK12 | BDNF | MKO3 | -0.12 | 1 | 0.79 |
| 43 | MAP3K1=MK12 | INS | HSPB1 | -0.12 | 1 | 0.79 |
| 44 | MAP3K1=MK12 | INS | MK12 | -0.12 | 1 | 0.79 |
| 45 | MAP3K1=MK12 | INS | MKO3 | -0.12 | 1 | 0.79 |
| 46 | MAP3K1=MK12 | EGCG | HSPB1 | -0.12 | 1 | 0.79 |
| 47 | MAP3K1=MK12 | EGCG | MK12 | -0.12 | 1 | 0.79 |
| 48 | MAP3K1=MK12 | EGCG | MKO3 | -0.12 | 1 | 0.79 |
| 49 | MAP3K1=MK12 | CONA | HSPB1 | -0.12 | 1 | 0.79 |
| 50 | MAP3K1=MK12 | CONA | MK12 | -0.12 | 1 | 0.79 |
| 51 | MAP3K1=MK12 | CONA | MKO3 | -0.12 | 1 | 0.79 |
| 52 | MAP3K1=MK12 | TNFA | HSPB1 | -0.12 | 1 | 0.79 |

| | | | | | | |
|---|---|---|---|---|---|---|
| 53 | MAP3K1=MK12 | TNFA | MK12 | -0.12 | 1 | 0.79 |
| 54 | MAP3K1=MK12 | TNFA | MKO3 | -0.12 | 1 | 0.79 |
| 55 | NACL=MK12 | NACL | HSPB1 | -0.16 | 0 | 0.17 |
| 56 | NACL=MK12 | NACL | MK12 | -0.16 | 0 | 0.17 |
| 57 | NACL=MK12 | NACL | MKO3 | -0.16 | 0 | 0.17 |
| 58 | SLP76=AKT1 | ANTICD3 | AKT1 | -0.14 | 1 | 0.46 |
| 59 | SLP76=AKT1 | ANTICD3 | GSK3A | -0.14 | 1 | 0.46 |
| 60 | SLP76=AKT1 | EGCG | AKT1 | -0.14 | 1 | 0.46 |
| 61 | SLP76=AKT1 | EGCG | GSK3A | -0.14 | 1 | 0.46 |
| 62 | ZAP70=GRB2 | ANTICD3 | HSPB1 | 0.00 | 0 | 0.33 |
| 63 | ZAP70=GRB2 | ANTICD3 | MK12 | 0.00 | 0 | 0.33 |
| 64 | ZAP70=GRB2 | ANTICD3 | MP2K1 | 0.00 | 0 | 0.33 |
| 65 | ZAP70=GRB2 | ANTICD3 | MKO3 | 0.00 | 0 | 0.33 |
| 66 | ZAP70=GRB2 | EGCG | HSPB1 | 0.00 | 0 | 0.33 |
| 67 | ZAP70=GRB2 | EGCG | MK12 | 0.00 | 0 | 0.33 |
| 68 | ZAP70=GRB2 | EGCG | MP2K1 | 0.00 | 0 | 0.33 |
| 69 | ZAP70=GRB2 | EGCG | MKO3 | 0.00 | 0 | 0.33 |
| 70 | ZAP70=SLP76 | ANTICD3 | AKT1 | 0.00 | 1 | 0.38 |
| 71 | ZAP70=SLP76 | ANTICD3 | GSK3A | 0.00 | 1 | 0.38 |
| 72 | ZAP70=SLP76 | EGCG | AKT1 | 0.00 | 1 | 0.38 |
| 73 | ZAP70=SLP76 | EGCG | GSK3A | 0.00 | 1 | 0.38 |
| | | | **NTZ** | | | |
| 1 | !ANTICD3=PAG | ANTICD3 | STAT5 | -0.06 | 1 | 0.42 |
| 2 | ANTICD3=WNK1 | ANTICD3 | WNK1 | 0.00 | 1 | 0.42 |
| 3 | BDNF=GRB2 | BDNF | MP2K1 | -0.05 | 1 | 0.47 |
| 4 | CONA=STAT1 | CONA | STAT1 | -0.06 | 1 | 0.42 |
| 5 | GP130=JAK1 | IL6 | STAT1 | -0.10 | 1 | 0.95 |
| 6 | MAP2K4=MK12 | TNFA | HSPB1 | -0.15 | 0 | 0.21 |
| 7 | MAP2K4=MK12 | TNFA | MK12 | -0.15 | 0 | 0.21 |
| 8 | MAP2K4=MK12 | TNFA | MKO3 | -0.15 | 0 | 0.21 |
| 9 | MAP2K4=MK12 | TNFA | STAT5 | -0.15 | 0 | 0.21 |
| 10 | MAP3K1=MAP2K4 | TNFA | HSPB1 | -0.08 | 0 | 0.21 |
| 11 | MAP3K1=MAP2K4 | TNFA | MK12 | -0.08 | 0 | 0.21 |
| 12 | MAP3K1=MAP2K4 | TNFA | MKO3 | -0.08 | 0 | 0.21 |
| 13 | MAP3K1=MAP2K4 | TNFA | STAT5 | -0.08 | 0 | 0.21 |
| 14 | NACL=MK12 | NACL | HSPB1 | -0.06 | 0 | 0.26 |
| 15 | NACL=MK12 | NACL | MK12 | -0.06 | 0 | 0.26 |
| 16 | NACL=MK12 | NACL | MKO3 | -0.06 | 0 | 0.26 |
| 17 | NACL=MK12 | NACL | STAT5 | -0.06 | 0 | 0.26 |
| 18 | REBIF=JAK1 | REBIF | STAT1 | -0.05 | 1 | 0.58 |
| 19 | TAK1=MK12 | IL1A | HSPB1 | -0.15 | 1 | 0.79 |
| 20 | TAK1=MK12 | IL1A | MK12 | -0.15 | 1 | 0.79 |
| 21 | TAK1=MK12 | IL1A | MKO3 | -0.15 | 1 | 0.79 |
| 22 | TAK1=MK12 | IL1A | STAT5 | -0.15 | 1 | 0.79 |
| 23 | TAK1=MK12 | LPS | HSPB1 | -0.15 | 1 | 0.79 |
| 24 | TAK1=MK12 | LPS | MK12 | -0.15 | 1 | 0.79 |
| 25 | TAK1=MK12 | LPS | MKO3 | -0.15 | 1 | 0.79 |
| 26 | TAK1=MK12 | LPS | STAT5 | -0.15 | 1 | 0.79 |
| 27 | TAK1=MK12 | POLYIC | HSPB1 | -0.15 | 1 | 0.79 |
| 28 | TAK1=MK12 | POLYIC | MK12 | -0.15 | 1 | 0.79 |
| 29 | TAK1=MK12 | POLYIC | MKO3 | -0.15 | 1 | 0.79 |
| 30 | TAK1=MK12 | POLYIC | STAT5 | -0.15 | 1 | 0.79 |
