## Supplementary Methods S1 for "Prediction of combination therapies based on topological modeling of the immune signaling network in Multiple Sclerosis"

***Subjects and clinical cohorts***

Patients fulfilled McDonald 2005 criteria (1) and disease subtype was defined using Lublin criteria (2). Patients were allowed to receive any therapy and should be stable in such therapy for the previous 6 months. Patients were recruited by their neurologist after signing informed consent. The study was approved by the IRB of each clinical center.

***xMAP assays development***

*1. Antibody and stimulus screening experiments for antibody selection and standardization*

Several antibodies against phosphorylated proteins involved in the PKN (from Biorad, Millipore and ProtAtOnce (PAO)) were screened to find antibody pairs with good readouts. The best pairs were selected using a computational optimization scheme, selecting pairs with a high signal-to-noise ratio (SNR) as indicated in previous work (3). The stim-screen experiment was used to identify good stimuli for PBMCs that activate at least one phosphoprotein and good assays that show at least 50% increase in at least one stimuli. The following initial signaling pathways were selected for the modeling and measuring of the phospho-signals: JAK/STAT, p38 MAPK, PI3K/AKT/mTOR, JNK MAPK, AP-1 MAPK, NFkB, translational control, protein tyrosine kinases, oxidative stress, and Vitamin D action. To define the list of xMAP assays, the following parameters were taken into account: 1) Relevance to the general pathways of interest in the context of MS; 2) Direct references to the gene/protein in the context of MS; 3) Expected changes in the *in vitro* response to MS drugs (IFNB, FTY, NTZ, GA, dimethyl-fumarate (DMF) or teriflunomide (Terif)); 4) References from GWAS study; 5) Relevance to the environmental factors related to MS risk (vitamin D).

Stimulus screen experiments were performed in 96-well plates. The 96 well of each plate were distributed among 1 male and 2 female samples (32 wells each) and each subject wells were stimulated with 28 stimuli. We used one plate for each time for sample collection (0 – 5 – 25 min after simulation), and therefore each subject has three plates. Thus, at the end of the stimulus screen experiments 32 lysates were obtained from the 28 different treatments tested (pooled 3 donors & 2 time points), and then each lysate was split into 5 in order to compare the results when using different multiplex assays: PAO_PLEX-1, PAO_PLEX-2, BIORAD_PLEX, MILLIPORE_PLEX, IDIBAPS_PLEX. The final pairs were selected using the two final optimal multiplexed antibody combinations: PAO_PLEX-1, PAO_PLEX-2. In these assays many proteins are activated and there is good consensus between the PAO_Plex assays with

BioRad_Plex and Millipore_Plex assays (although Millipore assays have consistently lower signals). For this reason, both PAO Plex and BioRad assays were used. The final list comprised 17 proteins of interest: AKT1, CREB1, FAK1, GSK3A, HSPB1, IKBA, JUN, MK03, MK12, MP2K1, PTN11, STA5A, STAT1, STAT3, STAT6, TF65, WNK1.

*3. Quality control*

For phosphoproteomic measurements, Protein A was used as a positive control (as a control of whether the stimulus was pipetted to the right well and to confirm the spiker was in the correct alignment when the stimulus was added). If a mismatch is detected between the apparent and the expected location of the protein A, the whole plate is discarded due to erroneous plate layout. Donor readouts above a consensus ranking were removed from the study.

*4. Bead Count Filtering*

Every reported measurement of the Luminex machine is the median of a distribution of beads. If the number of beads sampled were small (< 15) then the measurement was discarded as not statistically robust. This process filtered out 0.3% of the whole dataset. In addition, we also checked for "irregular" distribution (e.g. bimodal distribution) in the dataset.

*4. Biological noise management*

In order to control for biological noise, we performed the following checks: 1) Replicates: To determine the baseline cell levels, one experiment was performed in triplicate to obtain the coefficient of variation (CV) for each triplicate. There were 3 patients with high CV values and big discrepancies in the same plate that were therefore removed. 2) Negative control beads: bovine serum album (BSA) coated beads were used as a negative control for non-specific binding, which determines the background noise. As positive controls, R-Phycoerythrin (PE) coated beads were used to check for instrument errors.

*5. Identification of outliers (wells and patients)*

The phosphoprotein data was also curated to exclude single well errors that can distort the data. In order to identify outlying wells and patients, all the raw measurements were visually inspected. During this visual inspection, it was observed that there were some wells that were different dramatically across all the measured proteins from the rest of the dataset. Taking into account that changes of this magnitude can hardly be attributed to biological factors, it was decided to exclude these wells from the final dataset as outliers as explained below. Patients who were "enriched" in these outlying wells were also excluded from the dataset. The identification of outlying wells took place as follow: First, it was computed the robust z-score of every "feature" (i.e signal upon stimulation with a specific compound) in the dataset across all patients. Then, the median z-score of every well was checked, and if it was above 4, meaning that more than half of the readouts were extremely high, the well was removed. After this initial screening, all patients' measurements were visually inspected individually to identify other types of irregularities that could have escaped the automated process. This process filtered out a total of 7 patients' samples and 37 wells.

*6. Analysis of phosphoprotein dataset*

After analyzing the initial phosphoproteomic dataset obtained from the first set of donor samples and performing the QC checkpoint described above, we applied the following pipeline:

- Quality control (QC): after application of the QC explained above, only 5% of the donors were removed. After data filtering, 82.8% of the 255 donor samples received were approved for the analysis.
- Phosphoproteomics measurements: The samples from 17 patients and 2 control were analyzed using the PAO multiplex assays. A 50% increase in the signal was defined as a positive response. We observed a food good quality of data (fold increase) and biological noise within the expected range in this dataset.

*7. Assessment of phospho assays in PBMCs from healthy controls*

Then, we tested the performance of PAO-PLEX assays in 3 healthy controls in order to identify the optimal stimuli of the 32 finally tested that activate the phosphoproteome network, and to identify donor-to-donor variability in healthy volunteers. We observed some donor-to-donor variability in the baseline levels although results are qualitatively similar (direction of the response after a given stimuli). After analyzing the stimuli used and their corresponding phosphoprotein (strong, weak or not signal), 20 stimuli were identified that provided consistent responses. This included pro-inflammatory or pro-oxidant stimuli (anti-CD3 (OKT3), concanavalin A, $H_2O_2$, IFN-gamma, IL1A, IL6, LPS, NaCl, PolyIC, TNF-alpha), inmunomodulatory stimuli (IFN-beta1a, S1P, vitamin D3) neuroprotectants or anti-oxidants (BDNF, EGCG, INS) and disease modifying drugs from MS (DMF, FTY and Teriflunomide) (**Supplementary Table S5**).

We performed two levels of normalization:

1) Normalization of phosphorylation values ("phospho level"): To normalize phosphoproteomic datasets in order to perform protein-to-protein and condition-to-condition comparisons, the change in phosphorylation for each protein and each patient was calculated with respect to the control conditions. The significance of the differential phosphorylation of each protein with respect to the protein's mean was used to determine the degree of correction needed to perform the analysis.

3) Statistical normalization by logarithmic transformation (respect to the baseline levels ("response value")).

Since proteins phosphorylation was tested in many conditions (stimuli and culture media as control), the value of phosphorylation response to the stimulus in respect to the baseline level for each protein in each sample was defined as:

log2 (phospho_level_in_response_to_the_stimulus / phospho_level_in_medium)

These values were normally distributed and for this reason they were analyzed with parametric test (T test) for group comparison.

#### *Data normalization for logic modeling*

After signal reading, data was normalized. As described above, data were acquired at 0, 5 and 25 minutes after perturbation. The maximum between 5 and 25 measurements was selected to allow capturing signal transduction including late effects of network motifs such as negative feedback loops. Boolean logic modeling allows simulation for each node in the network as active or inactive. For some perturbations, signals were measured as being activated, inhibited or stay at the same level of phosphorylation as the control. To allow Boolean modeling of this three-state scenario, differential phosphorylation was calculated and the significant state chosen, thereby discarding the non-significant state and yielding the two most biologically relevant states. This was achieved by testing the log2 Fold Changes (FC) and selecting significant FCs to calculate the signaling activity for each individual interaction in a given patient. Measurements for the same protein that diverged from the significant state were replaced by NA to avoid biasing optimization. Finally, to avoid bias by outliers, a Hill-based non-linear normalization was applied. The normalization algorithm extends from (Saez-Rodriguez et al. 2009) and is shown in **Supplementary Figure S2** and the final dataset with all transformations in **Figure 2**.

#### *Modeling*

*1.Model complexity reduction*

Considering that by expanding the logic gates to all possible combinations of 2 or 3 inputs for each AND gate led to an underdetermined modeling, to avoid this, we assessed whether complexes were found between the proteins in the identifiable model. Using an iterative analysis via Bioservices (4) and the EBI's complex portal (5) (https://www.ebi.ac.uk/intact/complex), we determined that only the NFkB-IKB complex was present, therefore only this motif was included in the topology. Hence, based on molecular knowledge the model complexity was reduced to a degree that could be solved by model optimization.

*2. Optimization*

For each donor, 10 successful optimization runs were performed with a genetic algorithm set to 10 hours, a maximum of 100,000 generations, a population size of 100 individuals, an elitism of 2 individuals propagated through iterations, and a relative tolerance compared to error of the best solution of 0.00005 a.u. To assess the validity and robustness of our modeling approach, model quality was examined as shown in **Supplementary Figure S3**. After merging all model solutions into a single, robust model for each donor, the resulting donor-specific models were grouped according to treatment and disease subtype. Therefore, the signaling activity calculated for each individual interaction in a given donor was mean averaged with the signaling activities of all other donors within the same group for that same interaction. All presented analyses have been performed with R 3.4.1 and the RWTH computing cluster, running an LSF job scheduling system, version 9.1.3.0. The CellNOptR package was used in version 1.22.0.

*3. Differential activation for subgroup networks*

To calculate differential activation of signaling interactions between subgroup models, we (i) first filtered out inactive interactions by selecting those interactions whose signaling activity was below the respective upper quartile in at least one of the subgroup models. Then we (ii) identified subgroup differences by subtracting one model from the other. Finally, (iii) we identified the differentially active interactions in analogy to the co-druggability score (see **Methods**): Those interactions whose difference between the two groups was in the upper quartile of the absolute differences were assigned to the respective single group only. Those interactions not passing this filter, i.e. with a difference close to zero are considered to be active in both subgroups.

### Supplementary References


1. Polman CH, et al. (2005) Diagnostic criteria for multiple sclerosis: 2005 revisions to the "McDonald Criteria." *Ann Neurol* 58(6):840–846.

2. Lublin FD, et al. (2014) Defining the clinical course of multiple sclerosis: the 2013 revisions. *Neurology* 83(3):278–286.

3. Poussin C, et al. (2014) The species translation challenge-a systems biology perspective on human and rat bronchial epithelial cells. *Sci Data* 1:140009.

4. Cokelaer T, Pultz D, Harder LM, Serra-Musach J, Saez-Rodriguez J (2013) BioServices: a

common Python package to access biological Web Services programmatically. *Bioinformatics* 29(24):3241–3242.

5. Meldal BHM, et al. (2015) The complex portal--an encyclopaedia of macromolecular complexes. *Nucleic Acids Res* 43(Database issue):D479–84.
